# VACmap: An Accurate Long-Read Aligner for Unraveling Complex Genomic Rearrangements

**DOI:** 10.1101/2023.08.03.551566

**Authors:** Hongyu Ding, Fritz J Sedlazeck, Christos Proukakis, Caoimhe Morley, Marco Toffoli, Anthony HV Schapira, Zhirui Liao, Lianrong Pu, Shanfeng Zhu

## Abstract

Inversions, duplications, and other critical medically challenging variations are often ignored by routine genetic analyses. This is due to the complexity of these alleles but also because of the inability to accurately align them with state-of-the-art methods. We introduce VACmap, a novel non-linear long-read mapping method designed to improve detection of these difficult genomic regions, including critical genes like *LPA* and *GBA1*, which are significant risk factors for cardiovascular disease and Parkinson’s disease, respectively.

## Main

Structural variations (SVs) have emerged as crucial elements in genomic research, with their biological significance becoming increasingly apparent [1-4]. These genomic alterations, typically defined as changes of 50 base pairs or more, encompass five main categories: insertions, inversions, deletions, duplications, and translocations, as well as combinations thereof [1]. The impact of SVs on genomic composition is substantial, affecting more nucleotides than any other type of genetic variation [3, 5-7]. Consequently, their influence is being recognized across evolutionary processes, human health, and contribute to both Mendelian and complex diseases as well as cancer development [8-11]. Despite their critical importance, our understanding of SVs remains limited [12]. This is in part due to the complexities of complex SV but also the inability of the current state of art aligners to represent them. Research has predominantly focused on simpler SV classes, such as deletions and insertions, while more intricate variations—like duplications, inversions, and complex clustered events—are often misrepresented and thus missed. This gap in our analytical approach hinders a comprehensive understanding of these important genomic phenomena [13, 14], despite their importance being highlighted in several publications already [7-11, 14-16].

These publications have mainly been driven by long-read sequencing that indeed enables the characterization of tandem repeats and thus regions where SV are predominantly observed [17]. Despite their advancements, long-reads are challenging to align due to their length and generally higher sequencing errors. In the past, others have highlighted that specialized aligners are required to accurately align them using predominantly linear alignments to the reference where multiple linear subalignments represent a potential SV. These sub-alignments are formed by identifying small, exact matching subsequences (k-mers) between the read and the reference sequence. Linear alignment algorithms then attempt to extend these matches in both directions until significant differences between the read and reference are detected. However, this process splits the read into segments and results in a large number of redundant subalignments, particularly in repetitive regions. In these regions, each segment of the read may map to multiple locations in the reference, making it difficult to determine the correct alignment. To improve this, long-read mapping methods have introduced additional steps aimed at choosing the most suitable subset of alignments to represent the complex structures introduced by SV. For example, minimap2 employs a greedy approach, selecting the highest-scoring sub-alignment at each step until all alignable regions of the read are accounted for [18]. In contrast, YAHA uses a graph-based algorithm that optimizes alignment selection by considering factors such as alignment length, quality, and genomic position [19]. NGMLR takes a similar approach but enhances the process with a scoring function and dynamic programming to identify the optimal combination of sub-alignments with the highest overall score [20]. Despite these advancements, post-alignment processes for determining the ideal set of sub-alignments are often inadequate because of the complexity of the allele and the underlying repetitive regions. They fail especially for duplication, inversion, and translocation, which are often missed or falsely identified by the linear alignment algorithms. Duplications are often misaligned as insertion because linear alignment algorithms prefer a single continuous alignment and treat duplication, which would require splitting reads, as insertion. Similarly, inversion and translocation, are often rather misaligned as splitting the reads is penalized and thus avoided.

To overcome this limitation of current methods, we propose VACmap, a long-read mapping method specially designed to improve the representations of all SV with a focus on complex genomic rearrangements. VACmap eliminating the conversational necessity of the post-alignment step for alignment selection by allowing both linear and non-linear extension in the alignment process. This novel approach simplifies the conversational workflow for read mapping and significantly improves the representation of complex alleles, providing a more accurate and comprehensive view of SVs.

Figure 1a-d gives an overview of VACmap’s non-linear mapping approach. The most important differentiation between VACMap and other approaches is implemented after initial matches between reference and read sequences have been identified. Here existing methods try to conserve the order of all subalignments by heavily penalizing splits when searching chains of matches maintained. The linear alignment approach can efficiently model genomic alterations such as deletions, insertions, and substitutions since these don’t break the collinearity of a chain. However, the linear approach penalizes the detection of complex SV such as duplication, inversion, translocation, or combinations of SV. In VACmap, we propose a novel hybrid alignment algorithm, which combines both linear and non-linear linkage approaches in a chain. In detail, VACmap represent matches as quadruples called ‘anchors’, which include the start positions in the long read and reference sequences, the strand match, and the anchor’s length. They are ordered by their start positions in the long read. The VACmap’s non-linear chaining algorithm then promotes the extension of the chain to subsequent anchors that preserve a strictly linear relationship with the preceding anchor, enhancing this connection with a positive score. Conversely, it penalizes extensions to anchors that disrupt this linearity by assigning negative scores to such connections. Then the optimal non-linear alignment of the entire sequence is the chain with the highest aggregate score (the longest path). Each of these linear subalignments can be extracted by dividing the non-linear alignment at the non-linear junction, eliminating the traditional necessity of additional post-alignment steps that reconstruct genomics rearrangement from a pool of error-prone independent subalignments. See methods for details.

**Figure 1.**
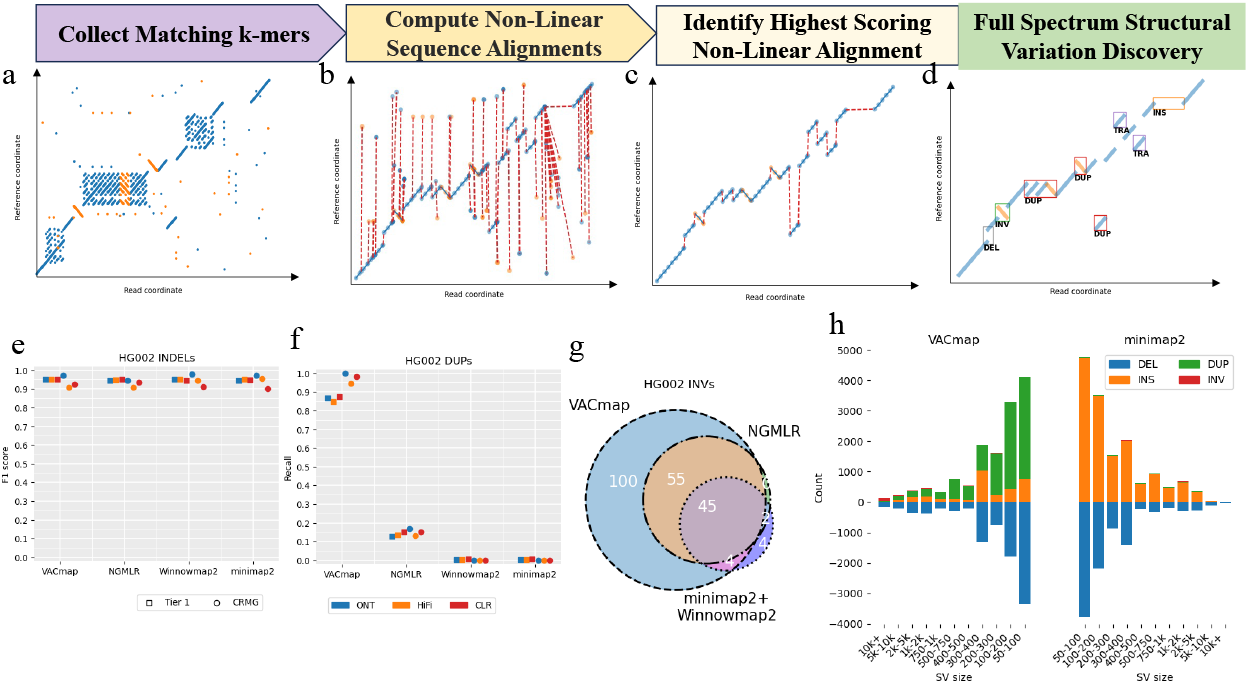
The workflow of VACmap non-linear alignment algorithm and impact on downstream SV detection. **a-d** The workflows of the VACmap approach for mapping complex rearrangements. **a**. Collect matching k-mers between the long read and reference (blue for the forward strand, orange for the reverse strand). **b**. Computing non-linear alignments. **c**. Extract the non-linear alignment with the highest score. **d**. Divide the highest score non-linear alignment into multiple linear subalignments and easy to interpret SV. **e, f**. Performance assessment of SVIM using four aligners’ alignments on GIAB benchmarks. **g**. Venn diagram showing the number of inversions discovered by SVIM call sets using alignments from four different mapping methods on HiFi data. **h**. The distribution of types and lengths among variants detected by SVIM on VACmap and minimap2 produced alignments.

We evaluated the SV detection performance of VACmap, NGMLR, Winnowmap2, and minimap2 alignments using the GIAB benchmark set [4, 21] with SVIM [22]. Truvari [23] was used to assess precision, recall, and F1 scores. Before evaluation, SVIM’s tandem duplication calls were relabeled as insertions to allow for comparability to GIAB assembly-derived benchmark. As expected all four alignment approaches demonstrated similar performance in detecting deletions and insertions in both GIAB tier 1 and CMRG regions (Figure 1e). And the runtime of VACmap is faster than NGMLR and Winnowmap2, but slower than minimap2. However, VACmap requires approximately half the memory usage compared to the other aligners (Supplementary Table 1). To evaluate SVIM’s sensitivity in detecting duplications using alignments from different tools, we isolated tandem duplication calls within the GIAB benchmark set using REPTYPE annotation. The results (Figure 1f) showed that SVIM, using VACmap-produced alignments, exhibited the highest sensitivity for duplication detection, identifying approximately 70% and 80% more duplications compared to other alignment approaches in the GIAB tier 1 and CRMG regions, respectively. This is highly important for the interpretability of the impact of SV. The SV distribution detected by SVIM using VACmap-produced alignments differed notably from other approaches (Figure 1h, Supplementary Figures 1). The VACmap approach suggested that multiple insertions are indeed duplications, aligning with previous findings [14]. The enhanced ability to accrue mapping duplication segment also enabled VACmap to better characterization of a previously reported de novo variation [24] (Supplementary Figures 2). This variation, located at chr14:23,280,711 (GRCh38 coordinate), was initially reported as a de novo insertion due to different insertion sizes in the child (HG002: 537 bp) and parents (HG003: 214 bp and HG004: 15 bp). With VACmap alignment, this de novo insertion was revealed to be a 109 bp Variable Number Tandem Repeat (VNTR) with different repeat number in the child (n=5) and paternal parent (n=2).

**Figure 2.**
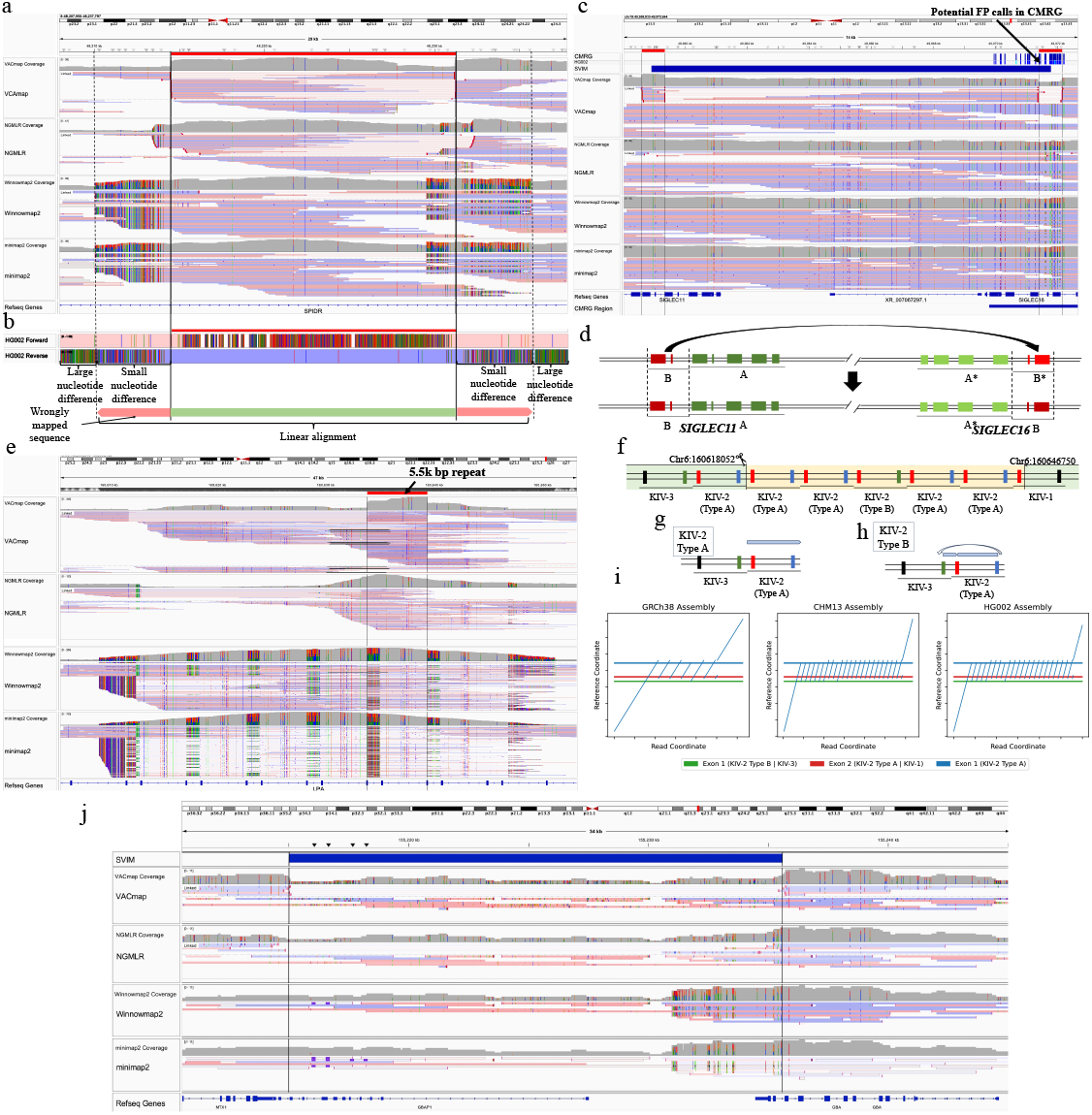
Comparison of four aligners on challenging gene regions. **a**, The IGV visualization of a segmental duplications flanked simple inversion at *SPIDR* gene locus. **b**. Comparison of sequences difference of HG002 assembly with GRCh37 assembly in inversion region at *SPIDR* gene locus. **c**. The IGV visualization of a potential *SIGLEC11* and *SIGLEC16* gene conversion event, the SVIM inversion prediction is shown in the top panel. **d**. Proposed scenario of gene conversions between *SIGLEC11* and *SIGLEC16* loci. **e**. The IGV visualization of alignments produced by four aligners in the *LPA* KIV-2 region. **f**. The illustration of GRCh38 reference modification and the exon structure of the KIV-2 domain. The exon 2 (red) in the type A KIV-2 repeat unit, type B KIV-2 repeat unit, and KIV-1 repeat unit have 100% identical sequences. The exon 1 (green) in the KIV-3 repeat unit and type B KIV-2 have 100% identical sequences. The light green and light orange regions indicate the reserved and removed regions, respectively. This figure is modified from [10]. **g, h**. The alignments scheme of type A KIV-2 sequence and type B KIV-2 sequence against modified GRCh38 reference. **i**. The dot plot depicting non-linear alignments generated by the VAC algorithm of GRCh38, CHM13, and HG002 assembly against modified GRCh38 reference. **j**. The IGV visualization of alignments produced by four aligners in the *GBA1 / GBAP1* region, with the SVIM deletion call shown at the top panel. Note, that Figures a, b, and c, d, e, j are under GRCh37, and GRCh38 coordinates, respectively.

Furthermore, the VACmap-SVIM callset included nearly all inversions (104/110) of the combined callset from minimap2, Winnowmap2, and NGMLR approaches and uncovered an additional 100 inversions (Figure 1g). For inversions that overlapped with previously reported inversion callset [25], the VACmap-SVIM pipeline identified nearly all inversions (49/50) reported by the other three approaches and identified an additional 12 inversions that were missed by the other three approaches (Supplementary Figures 3). We manually inspected the missed inversion from VACmap-SVIM pipeline and found out it’s actually an inversion flanked by inverted duplication and deletion. VACmap successfully resolves the complex structure but SVIM failed to detect it due to the intricacy structure not matching SVIM’s predefined rule to detect SV (Supplementary Figures 4).

Thus, highlighting that inversions remain challenging to resolve due to their locations often being surrounded by large segmental duplications. To further investigate this, we analyze the combined call set of 209 inversion regions from four aligners. Across all inversions, 32% (67/209) of them are overlapped with segmental duplications and half of them (39/67) are only detectable through VACmap alignment. For instance, VACmap alignment enables arcuate identification of a homozygous 16-kb inversion located in the *SPIDR* gene (Figure 2a), a gene is involved in DNA repair and associated with gonadal dysgenesis diseases [26]. On the contrary, other aligners’ alignments are less reliable, as they showed more mismatch bases (ie. signal of wrongly mapping of reads [20]) and inconstant breakpoints across different read alignments. It’s evidence in the size standard deviation of the inversion in SVIM calls is 254.2 and 2066.4 for VACmap and NGMLR alignment, respectively. A higher variance will be considered an unreliable SV prediction and assign a lower quality score (the SVIM quality score for this inversion is 13, 0 for VACmap, NGMLR alignment) and be discarded by downstream analyses. Figure 2b explains why NGMLR and other aligners failed to pinpoint the precise inversion breakpoints and force unrelate sequences to align together. The linear alignment approach will not terminate linear extension when reaching the inversion breakpoints since the subsequent sequence is too similar to the reference, leading to a wrongful linear alignment without the indication of inversion breakpoints. Supplementary Figures 5-9 showed additional examples of VACmap’s improved capabilities in mapping inversions.

However, SVIM may misinterpret complex inversion events as simple inversions. Figure 2c and Supplementary Figures 10 show an SVIM misidentified inversion is actually a potential *SIGLEC11* and *SIGLEC16* gene conversion event on the maternal haplotype. The *SIGLEC11* and *SIGLEC16* gene sequences are extremely similar in the region encoding the extracellular domain due to gene conversions [27]. The most recent gene conversion events were reported to occur one million years ago and involve regions A and A* in *SIGLEC11* and *SIGLEC16* genes, respectively [27] (Figure 2d). However, the VACmap alignment suggested a potential novel gene conversion event involving other similar regions B and B* in *SIGLEC11* and *SIGLEC16* genes, respectively (Figure 2d). This region B* in the *SIGLEC16* gene was previously observed by GIAB due to a dense series of heterozygous variants [21]. However, the gene conversion event is not resolved by GIAB due to limitations in the alignment technique and it causes dozens of false positive SNP calls in the GIAB CMRG benchmark set (Figure 2c). Further, VACmap alignment successfully resolved the homozygous *RHCE* and *RHD* gene conversion event when existing aligners failed and reduced over a hundred false positive SNP and indel calls in the GIAB CMRG benchmark set (Supplementary Figures 11-12).

We next assessed the *LPA* gene to highlight a medically important region that is further improved using VACmap. The complexity of this region raises due to high diversity in the population which represents 5-40 copies of the KIV-2 repeat in the *LPA* gene [10]. This copy number is inversely correlated with human lipoprotein(a) levels, which are strongly linked to coronary heart disease [10]. However, quantifying the KIV-2 copy number accurately poses significant challenges due to repetitiveness and thus the low mappability of sequencing reads in the *LPA* gene region [28]. We assessed the performance of four mapping methods by aligning PacBio HiFi and ONT sequencing data from human samples (CHM13 and HG002) against the GRCh38 reference genome. IGV visualizations revealed that NGMLR, Winnowmap2, and minimap2 produced alignments with excessive mismatches and less informative coverage information compared to VACmap (Figure 2e). VACmap demonstrated a superior ability to accurately represent KIV-2 repeats, showing clear and distinct coverage boundaries (Figure 2e, Supplementary Figures 13-15).

To simplify KIV-2 copy number determination, we modified the GRCh38 reference by removing the second to sixth KIV-2 repeat units and including the first 1000bp sequence of the follow-up KIV-1 unit (Figure 2f). We then realigned the PacBio HiFi and ONT data to the modified reference. Alignment dot plot and IGV visualizations indicated that VACmap-produced alignments (Figure 2i, Supplementary Figures 16-19) showed the expected alignment scheme of both type A and type B KIV-2 units (Figure 2g, 2h). Other mapping methods struggled to produce correct alignments despite the reduced complexity of the modified reference. Furthermore, the ONT reads facilitated the resolution of all 23 copies of the KIV-2 repeat unit in the CHM13 sample due to its longer read length compared to PacBio HiFi data (Supplementary Figures 20).

To further demonstrate the clinical utility of VACmap, we chose *GBA1*. This is a major risk factor for Parkinson’s disease [29], a challenging gene to analyze [30], which is prone to structural variants caused by recombination with a nearby highly homologous pseudogene (*GBAP1*). We previously detected using ONT long reads with adaptive sampling a pathogenic deletion which could not be correctly called after minimap2 or NGMLR alignment [31]. VACmap allowed SVIM to correctly report the breakpoints (Figure 2j), which is crucial in determining whether a deletion is pathogenic.

In summary, VACmap breaks through this long-standing barrier of inaccurate representing complex variants. This is achieved via an entirely new non-linear mapping approach and demonstrating the novelty and need of its method especially on inversions and other critical medically challenging genes such as *LPA* [21] and *GBA1*. Indeed, inversions remain challenging to resolve especially due to their location often surrounded by large segmental duplications [25]. Furthermore, these regions often form more complex events than simple inversions. Neither of forms complex or simple inversions are routinely detectable with state-of-the-art methods [25]. This is despite their clinical importance [32]. VACmap enables this detection with more precise alignments of read segments than any other method available due to its novel non-linear mapping approach. This further improves the characterization of complex duplications such as shown in KIV-2 a region in *LPA* itself and of gene / pseudogene recombination as shown in *GBA1*. VACmap can more precisely recapitulate the exact breakpoints within the reads which leads to an improved detection and thus will provide novel insights. These are only two examples of multiple medically important but challenging genes that VACmap can improve upon and thus deliver a more precise picture of the variants currently often missed by analytical methods [21].

## Method

### Variant-aware Chaining Algorithm

Current sequence alignment algorithms [13, 18-20, 33] attempt to align homology sequences via linear edits—deletions, insertions, and substitutions—that preserve the original sequence order and orientation. While effective for point mutations and small insertions or deletions (indels), this linear paradigm falls short when sequences exhibit complex genomic rearrangements such as duplications, inversions and translocations. Under this linear framework, duplications are often erroneously interpreted as insertions due to their identical signatures from a linear alignment perspective. Inversions, which do not result in a net gain or loss of genetic material, are challenging to distinguish from block substitutions. For more complex rearrangements—where multiple rearrangement events are clustered closely—the linear mapping approach attempts to reconstruct the complex structure by identifying all potential linear subalignments and selecting the most suitable subalignments from the pool of possibilities. The ideal representation, however, should consist of a series of linear subalignments connected by non-linear junctions at the breakpoints of these rearrangements. Yet, accurately retrieving the optimal set of subalignments is particularly challenging, especially for sequences located in highly repetitive regions or with substantial rearrangement complexity [34]. Moreover, since each linear subalignment is aligned independently, there is no assurance that a selection from locally optimal subalignments will culminate in a globally optimal alignment.

We present a novel solution with VACmap, a method that significantly deviates from the linear alignment model. VACmap integrates the Variant-aware Chaining (VAC) algorithm, which deftly combines linear and non-linear edits to achieve alignment. This dual-operation approach empowers VACmap to deliver a unified, globally optimal alignment that accurately reflects the genomic rearrangement in its entirety, bypassing the flawed post-alignment selection process.

To elaborate, our approach involved identifying k-mers that matched exactly between a reference sequence and a long-read sequence. These exact matches are expressed as quadruples, denoted as (x, y, s, k), and are termed anchors. Here, ‘x’ represents the start position of the anchor within the long-read sequence, while ‘y’ indicates the corresponding start position within the reference sequence. The variable ‘s’ signifies the orientation of the match, with a value of 1 indicating alignment with the forward strand and -1 indicating alignment with the reverse strand. The ‘k’ variable specifies the extent of the sequence encompassed by the anchor. Within the framework of the VAC algorithm, these anchors are conceptualized as nodes within a graph that is both directed and weighted. We apply specific rules to the linkage of these nodes: Only nodes that precede others in the long-read sequence can form connections to those that follow. To determine the weight of the directed edge from node[i] to node[j], where i is less than j, symbolized as w(i, j), we utilize a set of predefined formulas.

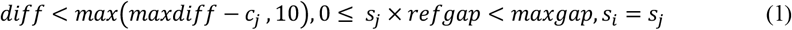

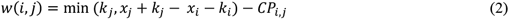

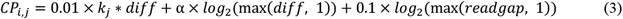

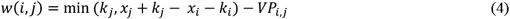

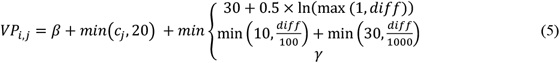

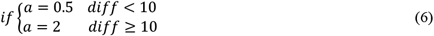

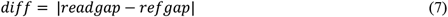

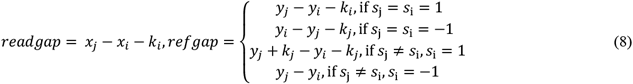

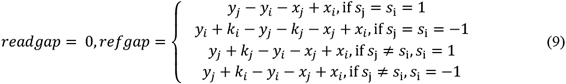

When two nodes exhibit strict collinearity, as defined by Formula 1, we calculate their connecting edge weight using Formula 2. Edges that fall under this category are termed “normal” edges. Formula 1 allows small deletions or insertions between nodes, which are regulated by the hyperparameter *maxdiff*, set by default to 50, and 30 for the first and second rounds of chaining, respectively. To incorporate an additional penalty for small indels, we adopt the affine-convex gap penalty model from minimap2 as outlined in Formula 3, which we refer to as the *CP* (colinear penalty). Conversely, when nodes do not conform to the conditions of Formula 1, we determine their edge weight with Formula 4, designating these as “variation” edges. The variation penalty, *VP*, is established through Formula 5, with its maximum value set to the sum of *β, γ*, and 20. The default settings for *β* and *γ* are 30 and 36, respectively. Given that k-mers occurring with high frequency are typically less informative [20], we impose an additional penalty, c[j], when extending a chain to a node associated with a frequently occurring k-mer, thus enforcing a stricter collinearity criterion. Here, c[j] represents the count of occurrences in the k-mer index corresponding to node j. For overlapping and non-overlapping anchors, we use formulas 8 and 9 to compute their distance, respectively. To further clarity regarding these principles, we provide illustrative examples in Supplementary Figure 21 to show the conditions for assigning normal and variation edges. The selection of hyperparameter values critically affects the quality of the non-linear alignment, with maxdiff and maxgap (default values of (50, 30) and 100, respectively) being particularly influential. A reduced maxdiff value tends to yield a more strictly collinear subalignment, while a smaller maxgap value helps to prevent the erroneous alignment of unrelated sequences.

To find the optimal non-linear alignment path among *N* nodes can be computed in *O(hN)* time if by dynamic programming. For a given *node[j]*, we can determine its maximum score *S(j)* and its best predecessor *node[i]* by utilizing the following formulas:

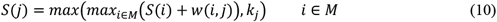

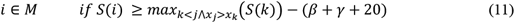

The *node[j]* best predecessor score *S*(*i*) has a low bound (Formula 10, 11). The low bound is equal to the current highest score *S[k]* minus (*β* + *γ* + 20), and *h* is equal to the size of set M.

Once the maximum score of all nodes is computed, we can identify the highest scores non-colinear chain by backtracking. The optimal set of colinear subalignments is recovered by discarding variation edges in the highest score chain.

### Local Index and Anchor Reduction

When sequence errors or SNPs occur in clusters, accurate mapping becomes challenging, especially when utilizing a large k-mer size setting and a minimizer k-mer sampling strategy. To mitigate this issue, VACmap adopts a similar approach used in a previous study [33] by constructing a local index with a smaller k-mer size, typically a 9-mer. The local index is built by collecting all possible k-mers from the previously computed highest-scoring chain’s covered reference genome region. Subsequently, all the k-mers in the long read are employed to query the index, obtaining matching information known as anchors. The highest scoring chain is then recomputed based on these anchors. However, the runtime of the VAC algorithm depends on the number of anchors.

Using a smaller k-mer size setting often results in an increased number of anchors, consequently escalating computational time. To address this issue, two strategies are employed to reduce the number of anchors and improve runtime efficiency. Firstly, VACmap iterates through the anchors and removes an anchor if the distance between the anchor and the previously computed highest-scoring chain exceeds a certain threshold (2000 bp). The distance between an anchor and the previous chain is defined as the distance between the anchor and its closest anchor in the chain under the reference sequence coordinate. Determining the closest anchor in the chain can be accomplished through a binary search, with a time complexity of *O(log*^*N*^), where N is the number of anchors. Secondly, VACmap merges two anchors *(x*_*i*_, *y*_*i*_, *s*_*i*_, *k*_*i*_) and *(x*_*j*_, *y*_*j*_, *s*_*j*_, *k*_*j*_) into a new anchor *(x*_*i*_, *min(y*_*i*_, *y*_*j*_*), s*_*i*_, *x*_*j*_ *+ k*_*j*_ *– x*_*i*_) if they satisfy certain conditions (Formula 12). The sorting of *N* anchors in ascending order based on their position in the long read allows the merging process to be computed in *O(N)* time using a hash table. The hash table utilizes an integer key and maintains a list of merged anchors as its value. For *anchor*_*i*_, if s_i_ equals 1, then the key is set to *y*_*i*_ *-x*_*i*_; otherwise, it is set to *-(y*_*i*_ *+ x*_*i*_). VACmap tests the last *anchor*_*j*_ in the list corresponding to a key. If the two anchors are overlapped or closely adjacent and the covered sequence in reference and long read is identical, VACmap updates the last anchor in the list; otherwise, *anchor*_*i*_ is appended to the list. If a key does not exist in the hash table, a new key-value pair is inserted. Finally, the merged anchors set can be obtained by traversing the hash table.

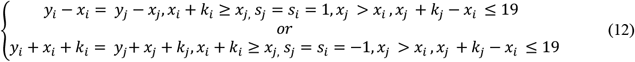

### Quality Control of Linear Subalignment

The presence of sequencing error and closely situated SNPs complicates the process of gathering sufficient anchors for the chaining process. To accommodate the occurrence of sequencing errors and clustered SNPs, VACmap employs a sizeable hyperparameter, maxgap, with a default setting of 100, within the VAC algorithm. However, a large maxgap value has the potential drawback of mistakenly aligning sequences that are not actually related. To address this problem, we assess the error ratio within each linear subalignment and exclude those that demonstrate a high propensity for errors. This error ratio is ascertained by employing edlib [35] to calculate the edit distance between the matched sequences in the reference genome and the long read. The edit distance obtained is then normalized by dividing the length of the shorter sequence involved in the comparison. A linear subalignment is deemed unreliable and is therefore rejected if its error ratio surpasses the predetermined threshold: 0.2 for sequences generated using PacBio CLR or ONT, and 0.1 for sequences generated using PacBio HiFi.

### Evaluation with GIAB callset

We evaluated four mapping methods, namely VACmap, NGMLR, Winnowmap2, and minimap2, using the GIAB Tier1 (v0.6) and CMRG (1.00) SV benchmark set for the HG002 human sample. These benchmark sets are available at the following links: https://ftp-trace.ncbi.nlm.nih.gov/ReferenceSamples/giab/release/AshkenazimTrio/HG002_NA24385_son/NIST_STRUCTURALVARIANT_v0.6/ and https://ftp-trace.ncbi.nlm.nih.gov/ReferenceSamples/giab/release/AshkenazimTrio/HG002_NA24385_son/CMRG_v1.00/, respectively. PacBio CLR, PacBio HiFi, and ONT sequencing data were accessible at s3://giab/data/AshkenazimTrio/. The SVIM (v1.4.2) and were used to detect SVs. Subsequently, we evaluated the SVs against the GIAB benchmark set using Truvari (v2.0.0). Command line parameters provided to these tools are listed in Supplementary Table 2.

## Data availability

This study utilized four human assembly genome sequences: GRCh37, GRCh38, CHM13 T2T (v2.0), and HG002 T2T (v1.1) [36]. They can be downloaded from the following links: http://ftp.1000genomes.ebi.ac.uk/vol1/ftp/technical/reference/phase2_reference_assembly_sequence/, https://ftp.1000genomes.ebi.ac.uk/vol1/ftp/technical/reference/GRCh38_reference_genome/, https://github.com/marbl/CHM13, and https://s3-us-west-2.amazonaws.com/human-pangenomics/T2T/HG002/assemblies/hg002v1.0.1.fasta.gz, respectively.

For the Pacbio CLR, Pacbio HiFi, and ONT sequencing data of the human sample HG002, HG003 and HG004. They can be downloaded from the following links: s3://giab/data/AshkenazimTrio, s3://giab/data/AshkenazimTrio, and s3://giab/data/AshkenazimTrio.

The Pacbio HiFi and ONT data for the human sample CHM13 were downloaded from SRA under accessions SRR11292120-SRR11292123 and s3://human-pangenomics/T2T/CHM13/nanopore/UW/chm13_UW_Guppy_3.6.0.fastq.gz, respectively.

The *GBA1* data are available from Prof C Proukakis on reasonable request, when compatible with consent/IRB restrictions.

## Code availability

VACmap are available on GitHub: https://github.com/micahvista/VACmap.

## Ethics statement

Ethics approval for the *GBA1* carrier was provided by the National Research Ethics Service London—Hampstead Ethics Committee as part of the RAPSODI study (www.rapsodi.com) [37]. Informed consent was provided.

## Acknowledgment

We would like to thank Dr. Kristoffer Sahlin (Stockholm University) and Dr. Mingfu Shao (Pennsylvania State University) for their helpful suggestions and insightful comments on the manuscript. This work has been supported by the National Natural Science Foundation of China (Grant No. 62272105), the Shanghai Municipal Science and Technology Major Project (Grant No. 2018SHZDZX01), 111 Project (Grant No. B18015), the ZJ Lab, the Shanghai Research Center for Brain Science and Brain-inspired Intelligence Technology. C.M. is supported by the MSA Trust.

## Author contributions

S.Z. supervised this study. H.D. designed the study and implemented the software. H.D., F.J.S., C.P. and C.M. performed the data analysis. H. D. drafted the paper. F.J.S., C.P., M.T., A.H.V.S., Z. L., L.P. and S.Z. modified the paper. All authors agree to the content of the final paper.

