## Supplementary data for "VACmap: An Accurate Long-Read Aligner for Unraveling Complex Genomic Rearrangements"

| **Dataset** | **Method** | **Run time(Hour)** | **Peak memory usage (GB)** |
| --- | --- | --- | --- |
| Pacbio CLR 69X | VACmap | 6.2 | **13.5** |
|  | NGMLR | 24 | 31 |
|  | Winnowmap2 | 7.5 | 33 |
|  | minimap2 | **3.3** | 29 |
| Pacbio HiFi 30X | VACmap | 4.2 | **11.6** |
|  | NGMLR | 12 | 27 |
|  | Winnowmap2 | 4 | 24 |
|  | minimap2 | **1** | 24 |
| ONT | VACmap | 5.8 | **14.7** |
|  | NGMLR | 32 | 34 |
|  | Winnowmap2 | 7.5 | 36 |
|  | minimap2 | **3.5** | 33 |

**Supplementary Table 1** **The runtime and peak memory usage of the four mapping methods.**

| **Tool** | **Purpose** | **Command** |
| --- | --- | --- |
| VACmap (v1.0) | Pacbio CLR or ONT read mapping  Pacbio CCS read mapping | vacmap -ref ref.fasta -read read.fasta -mode H -t 40 --markunbalancetra  vacmap -ref ref.fasta -read read.fasta -mode L -t 40 --markunbalancetra |
| minimap2 (v2.17) | Pacbio read mapping  ONT read mapping | minimap2 -t 40 -aYx map-pb ref.fasta read.fasta -o output.sam  minimap2 -t 40 -aYx map-ont ref.fasta read.fasta -o output.sam |
| winnowmap2 (v2.03) | Pacbio read mapping  ONT read mapping | winnowmap2 -t 40 -W repetitive k15.txt -aYx map-pb ref.fasta read.fasta -o output.sam  winnowmap2 -t 40 -W repetitive k15.txt -aYx map-ont ref.fasta read.fasta -o output.sam |
| NGMLR (v0.2.7) | Pacbio read mapping  ONT read mapping | ngmlr -t 40 -r ref.fasta -q read.fasta -o output.sam  ngmlr -t 40 -x ont -r ref.fasta -q read.fasta -o output.sam |
| SVIM (v1.4.2) | SV calling | svim alignment workdir sorted.bam ref.fasta --segment_overlap_tolerance 50 |
| Truvari (v2.0.0) | SV evaluation | truvari bench -b base.vcf -c comp.vcf -o workdir --passonly -p 0 -P 0 -r 500 --multimatch |

**Supplementary Table 2 Command line parameter used for this study.**

**a**

**b**

**c**

**d**

**e**

**f**

**g**

**h**

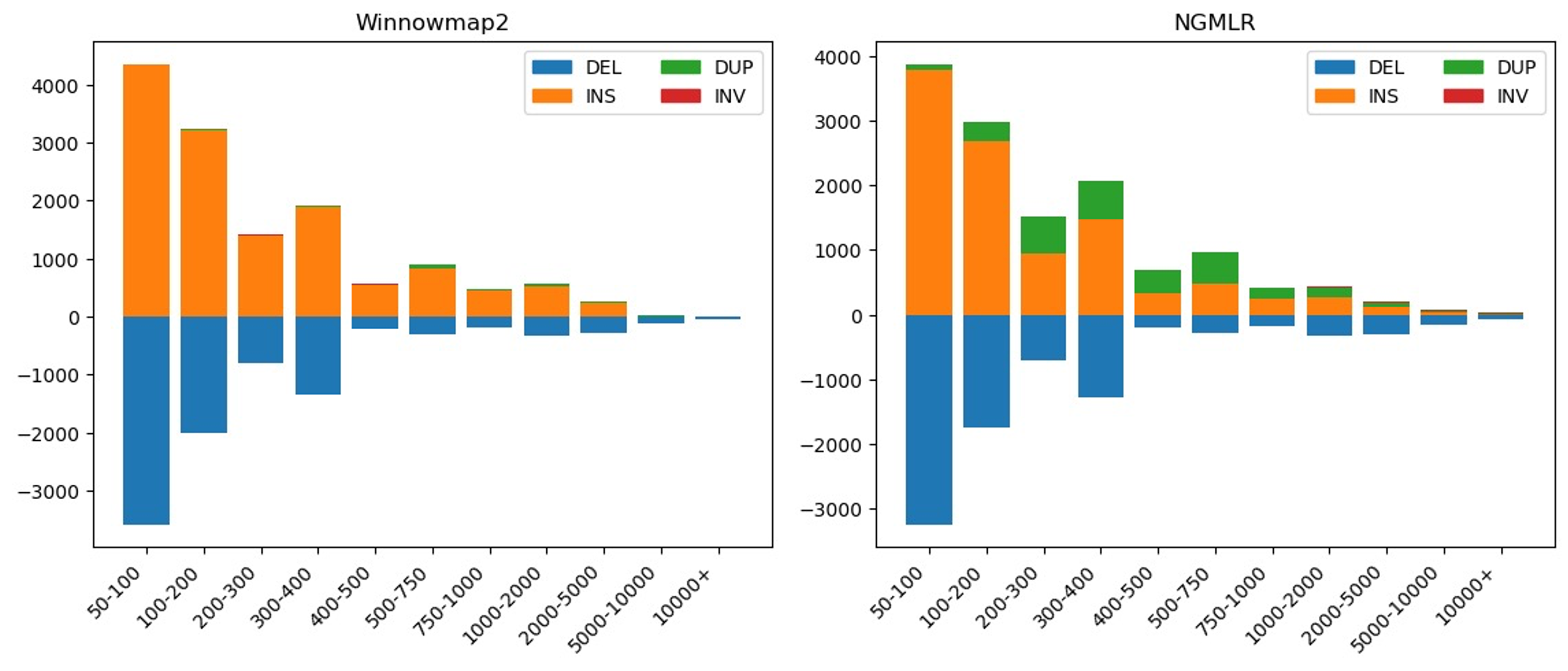

**Supplementary Figure 1 The distribution of types and lengths among variants detected by SVIM on Winnowmap2 and NGMLR produced alignments.**

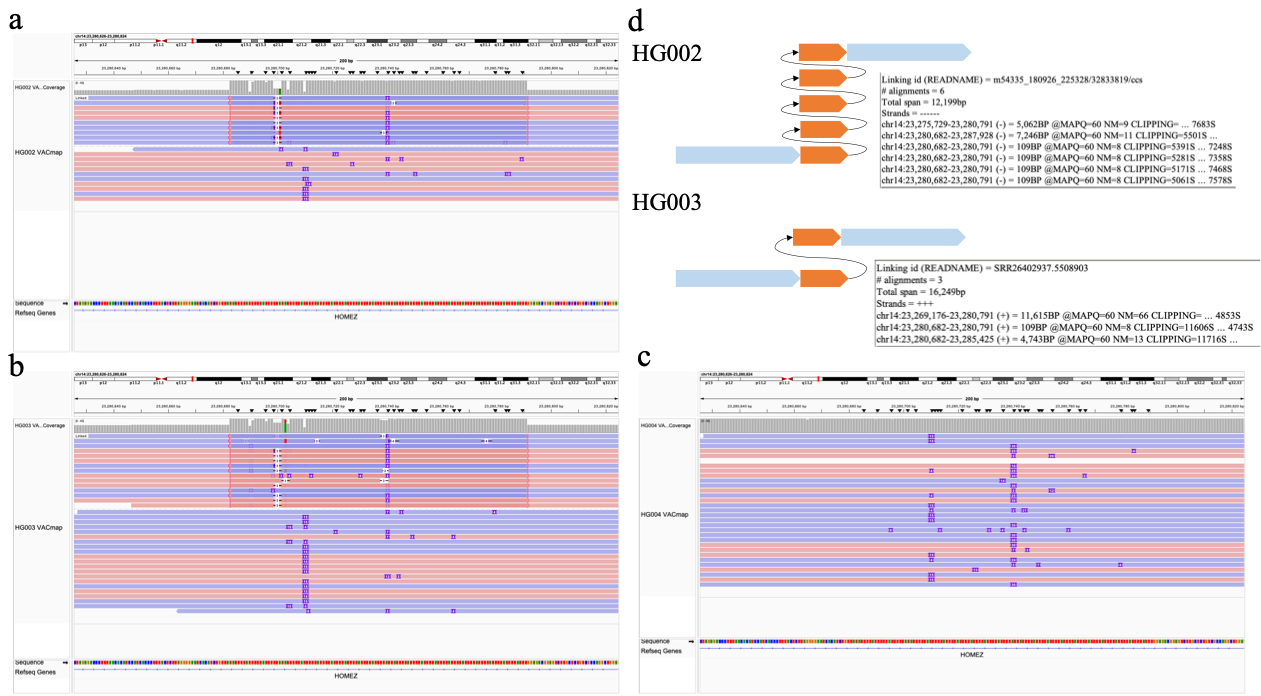

**Supplementary Figure 2** **Characterization of a de novo VNTR in HG002 at chr14:23,280,711 (GRCh38).** The VACmap alignments clearly show the de novo VNTR with different copy numbers in the child (HG002, n=5) and paternal parent (HG003, n=2) of the 109-bp repeat unit.

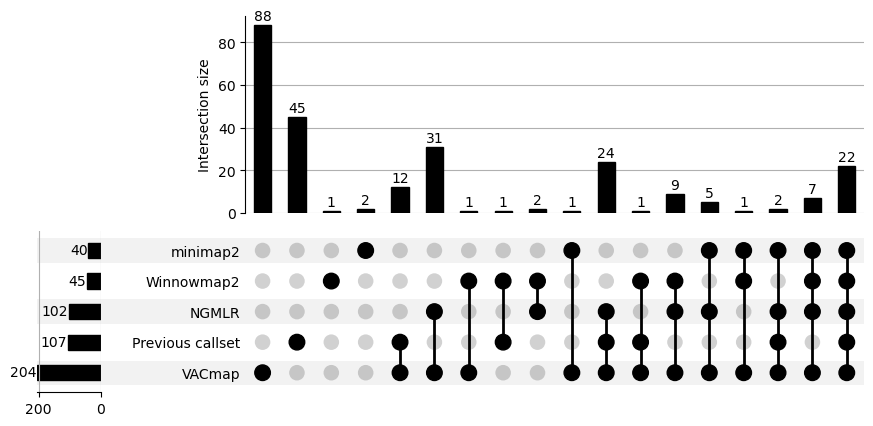

**Supplementary Figure 3 The upset plot of inversion detected by SVIM on VACmap, NGMLR, Winnowmap2 and minimap2 produced alignments and previous reported inversion callset.**

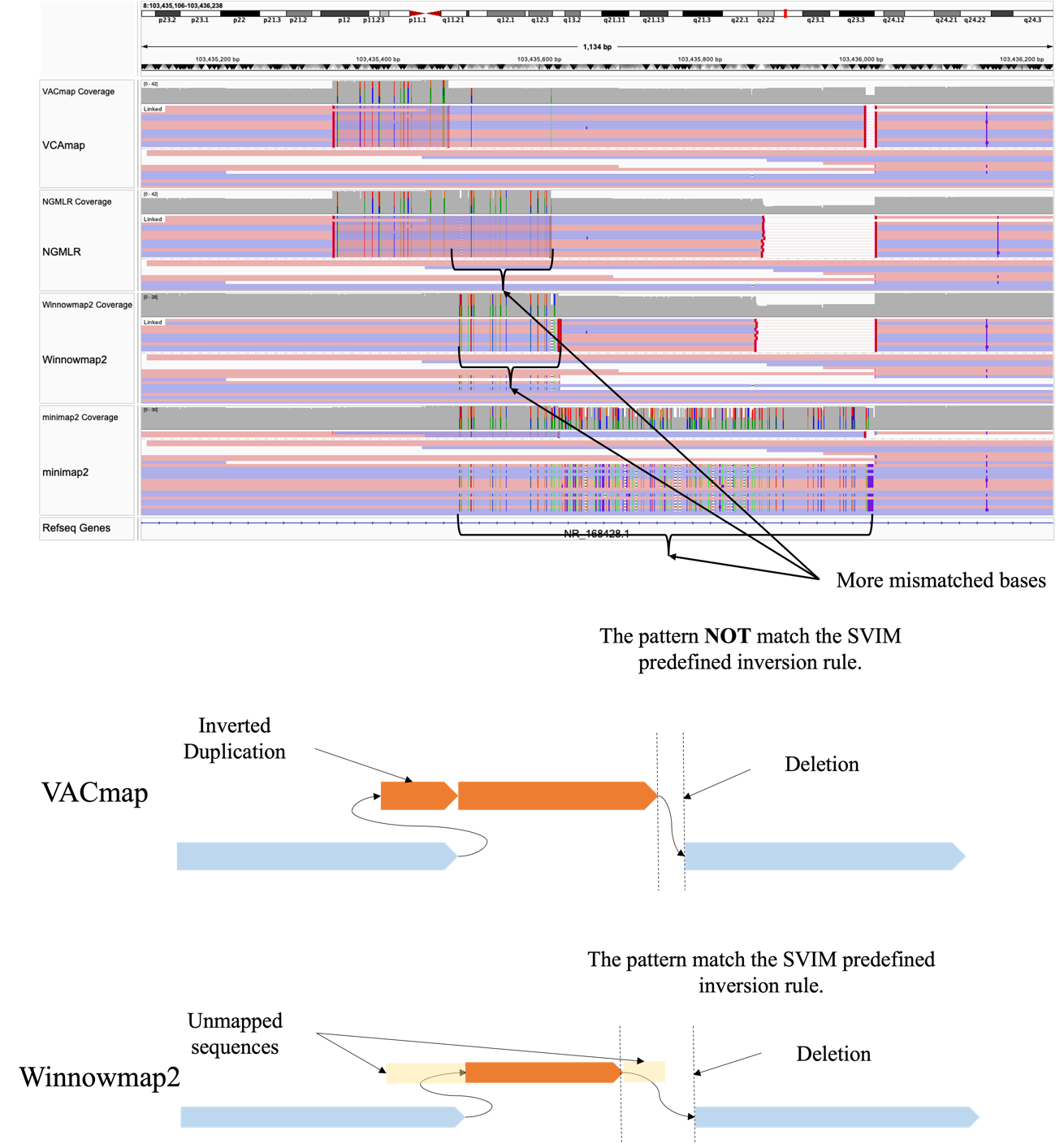

**Supplementary Figure 4 A inversion missed by SVIM due to the inversion complexity (GRCh37).** The VACmap-SVIM SV detection pipeline fails to identify this inversion because VACmap detected a more complex structure. VACmap's alignment suggests that the inversion is flanked by an inverted duplication and a deletion—an SV pattern that SVIM’s detection model is not designed to capture. This complex inversion pattern is also supported by NGMLR, though with less accuracy, as NGMLR's alignment indicates a higher number of mismatched bases compared to VACmap.

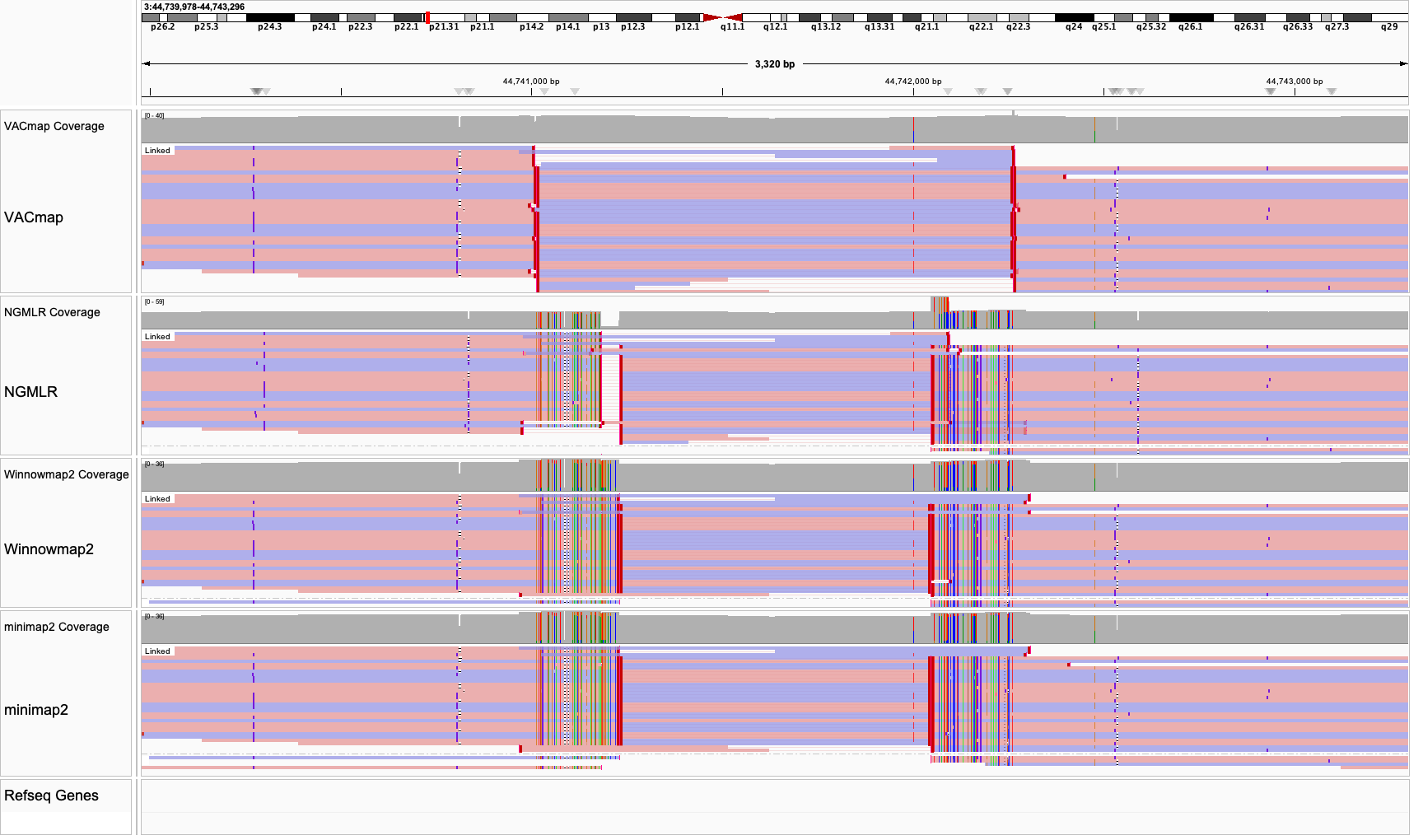

**Supplementary Figure 5 VACmap precisely identifies inversion breakpoints and reduces alignment mismatches (GRCh37).**

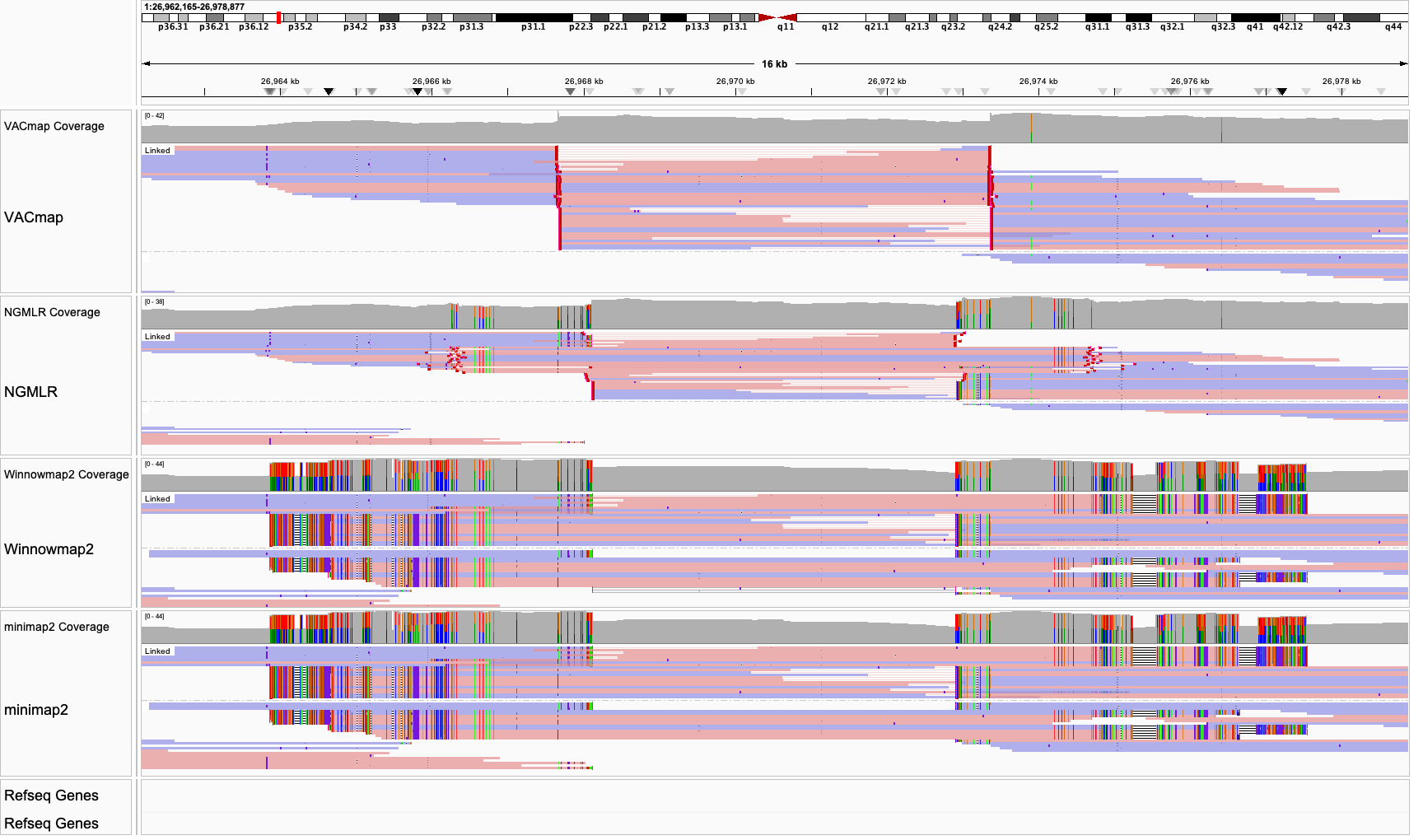

**Supplementary Figure 6 VACmap precisely identifies inversion breakpoints and reduces alignment mismatches (GRCh37).**

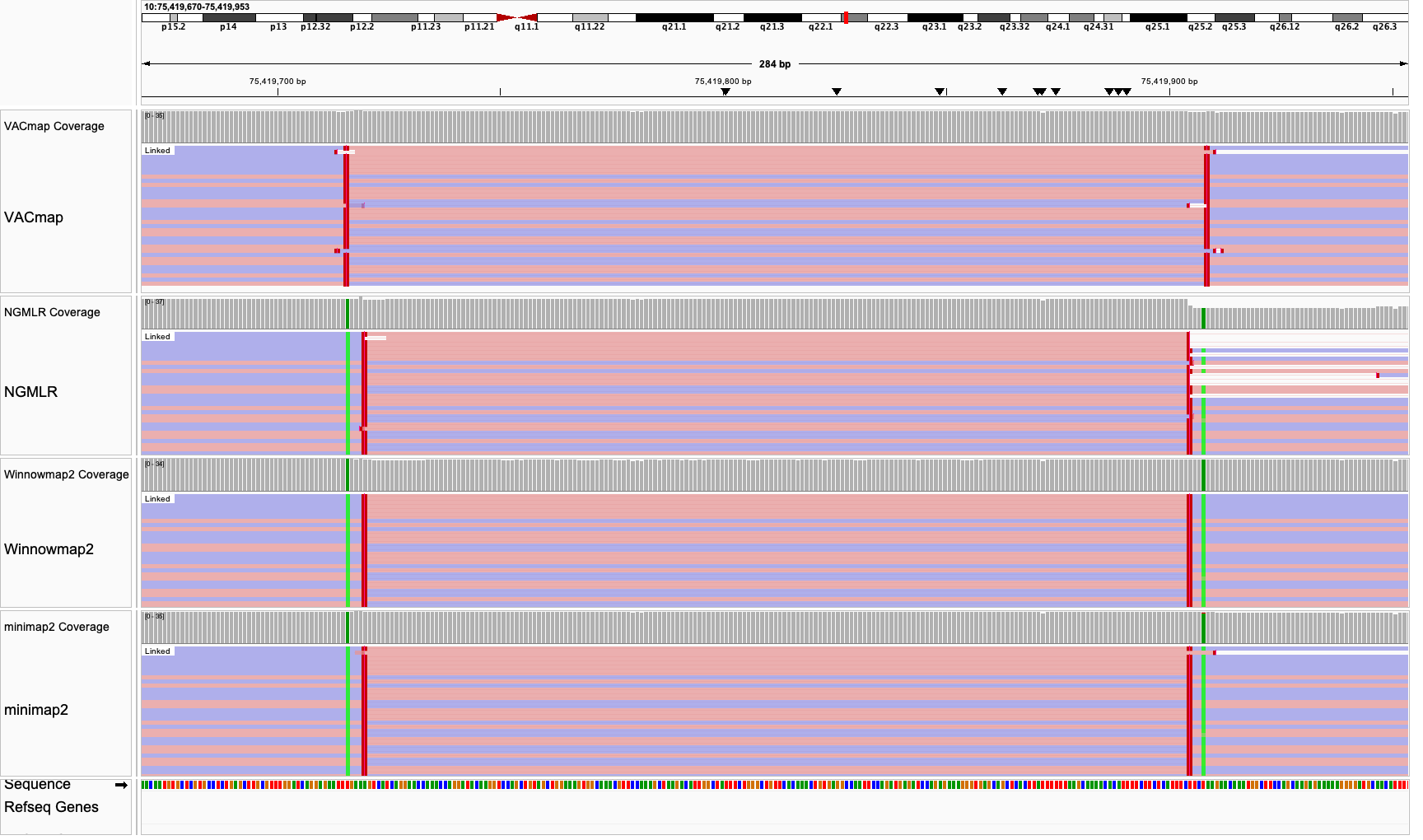

**Supplementary Figure 7 VACmap precisely identifies inversion breakpoints and reduces alignment mismatches (GRCh37).**

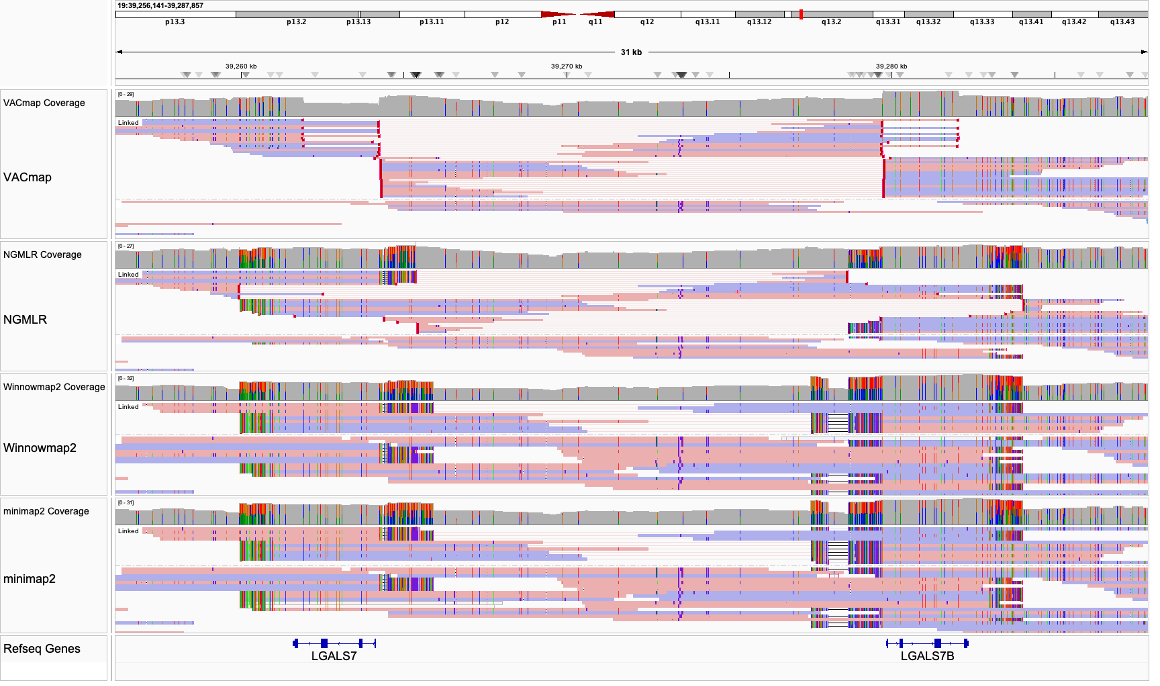

**Supplementary Figure 8 VACmap precisely identifies inversion breakpoints and reduces alignment mismatches (GRCh37).**

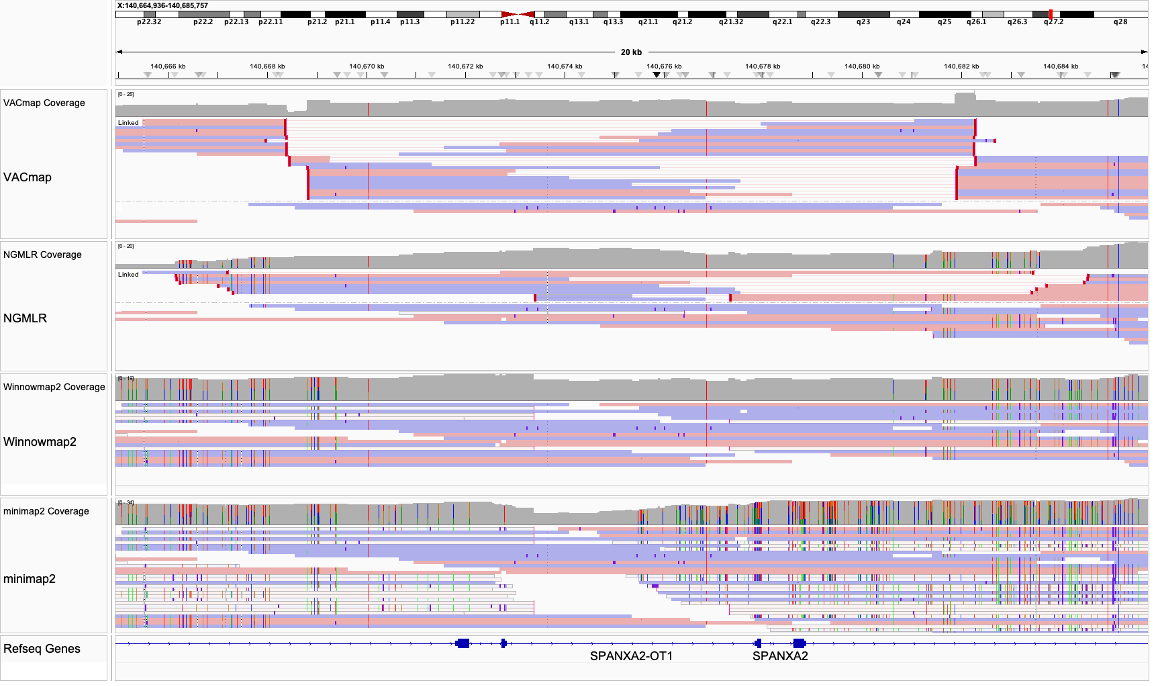

**Supplementary Figure 9 VACmap precisely identifies inversion breakpoints and reduces alignment mismatches (GRCh37).**

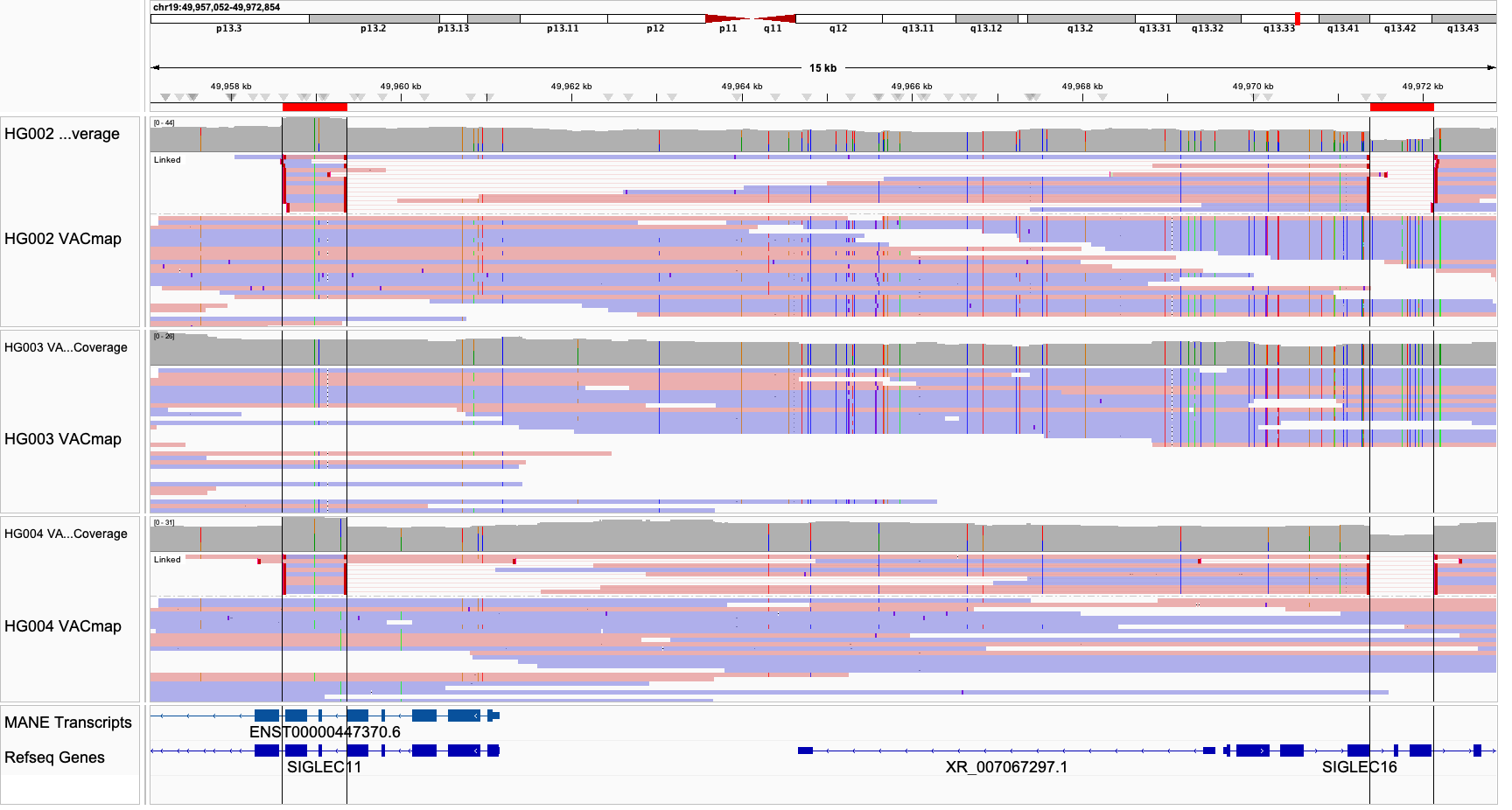

**Supplementary Figure 10 Potential *SIGLEC11* and *SIGLEC16* gene conversion event in the maternal haplotype of HG002 (GRCh38).**

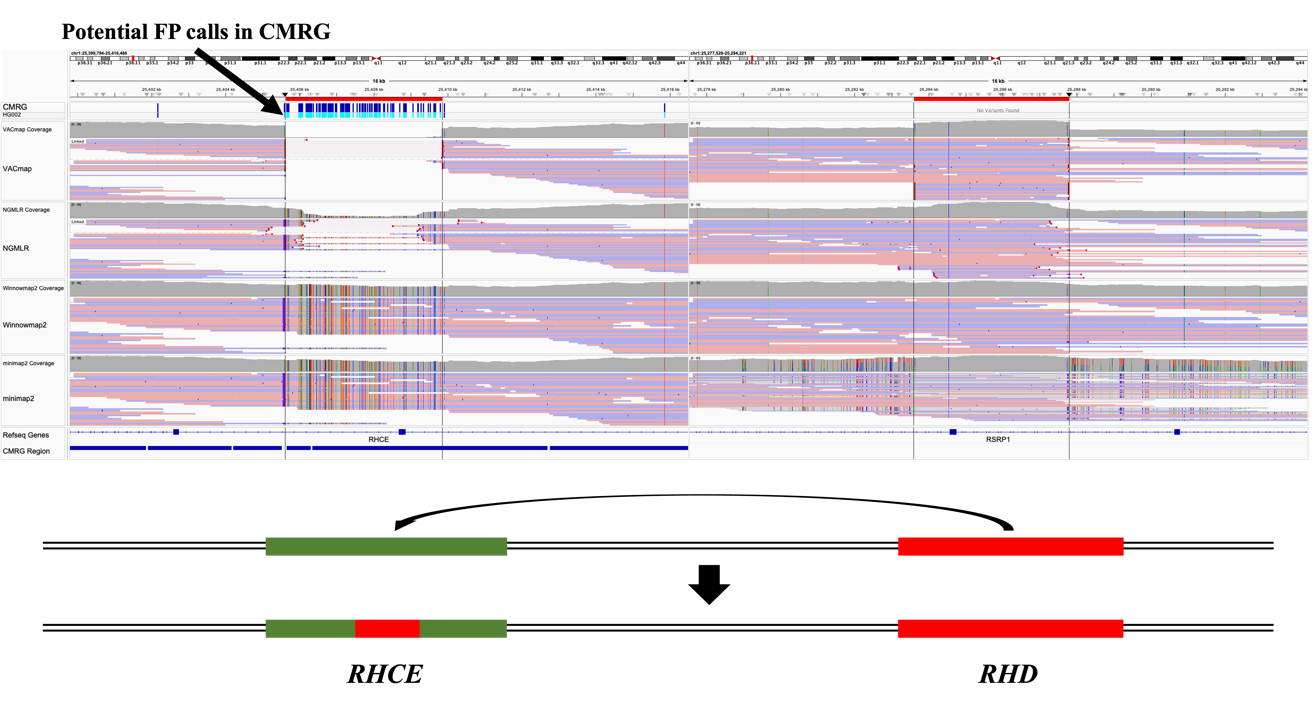

**Supplementary Figure 11 VACmap accurately identifies a gene conversion event between *RHCE* and *RHD* in HG002 (GRCh38).** This figure demonstrates VACmap's ability to precisely map a gene conversion event between the *RHCE* and *RHD* genes in HG002. The VACmap alignment clearly delineates the boundaries of the gene conversion region, which would be challenging to detect using standard alignment approaches.

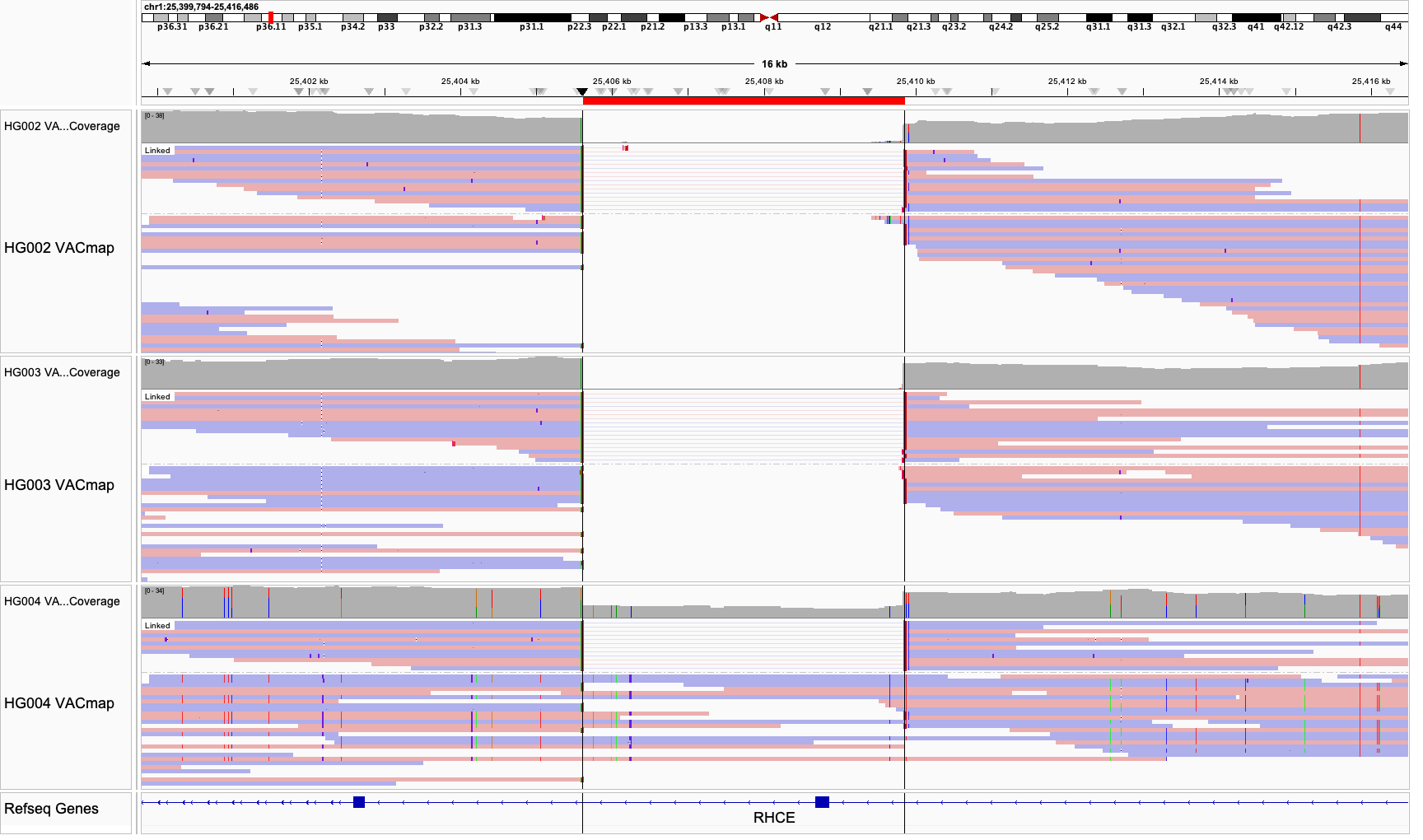

**Supplementary Figure 12 VACmap identifies *RHCE::RHD* gene conversion events in HG002, HG003, and HG004 (GRCh38).**

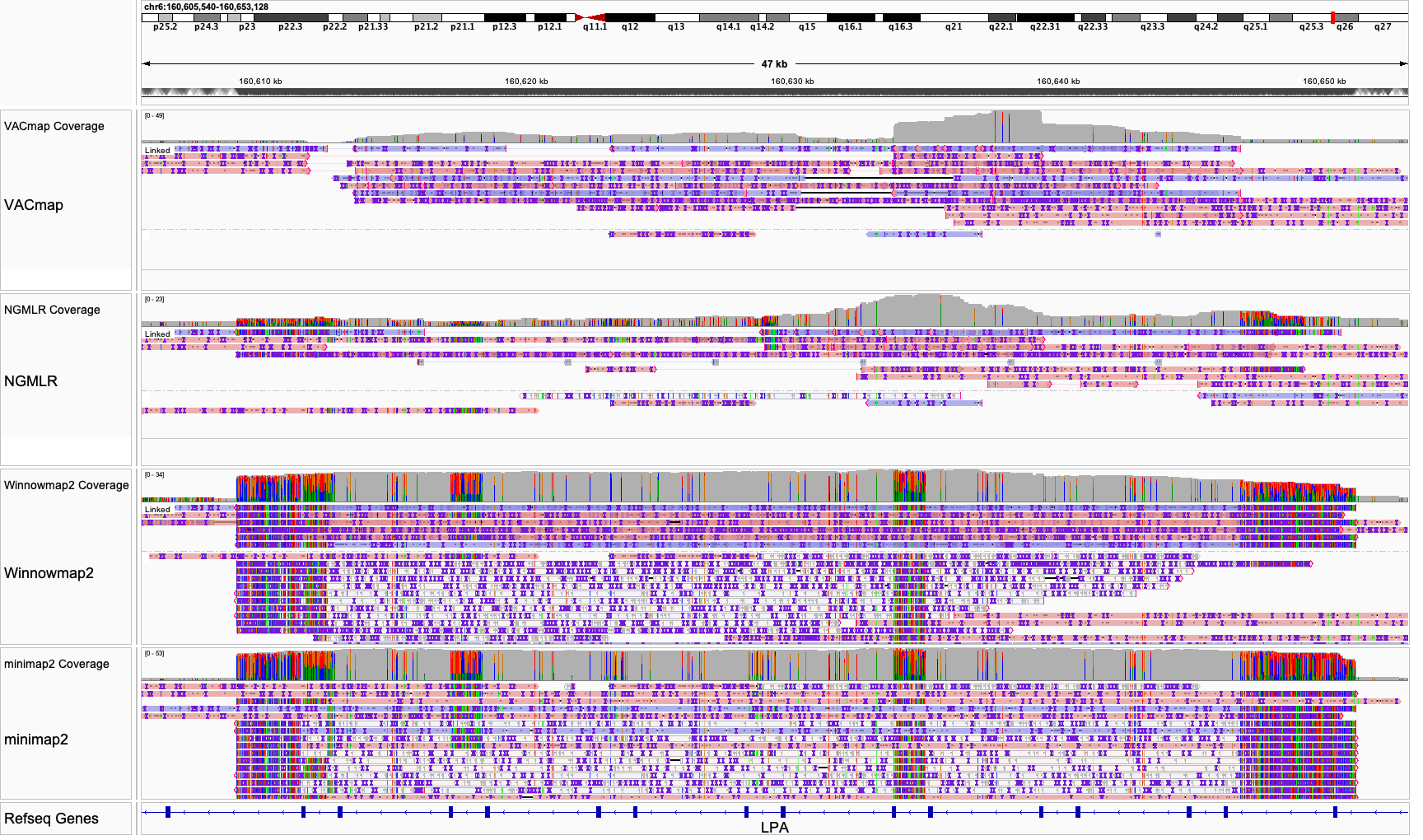

**Supplementary Figure 13** **The IGV visualization of CHM13 ONT alignments produced by four aligners under GRCh38 reference in the KIV-2 region** **(GRCh38).**

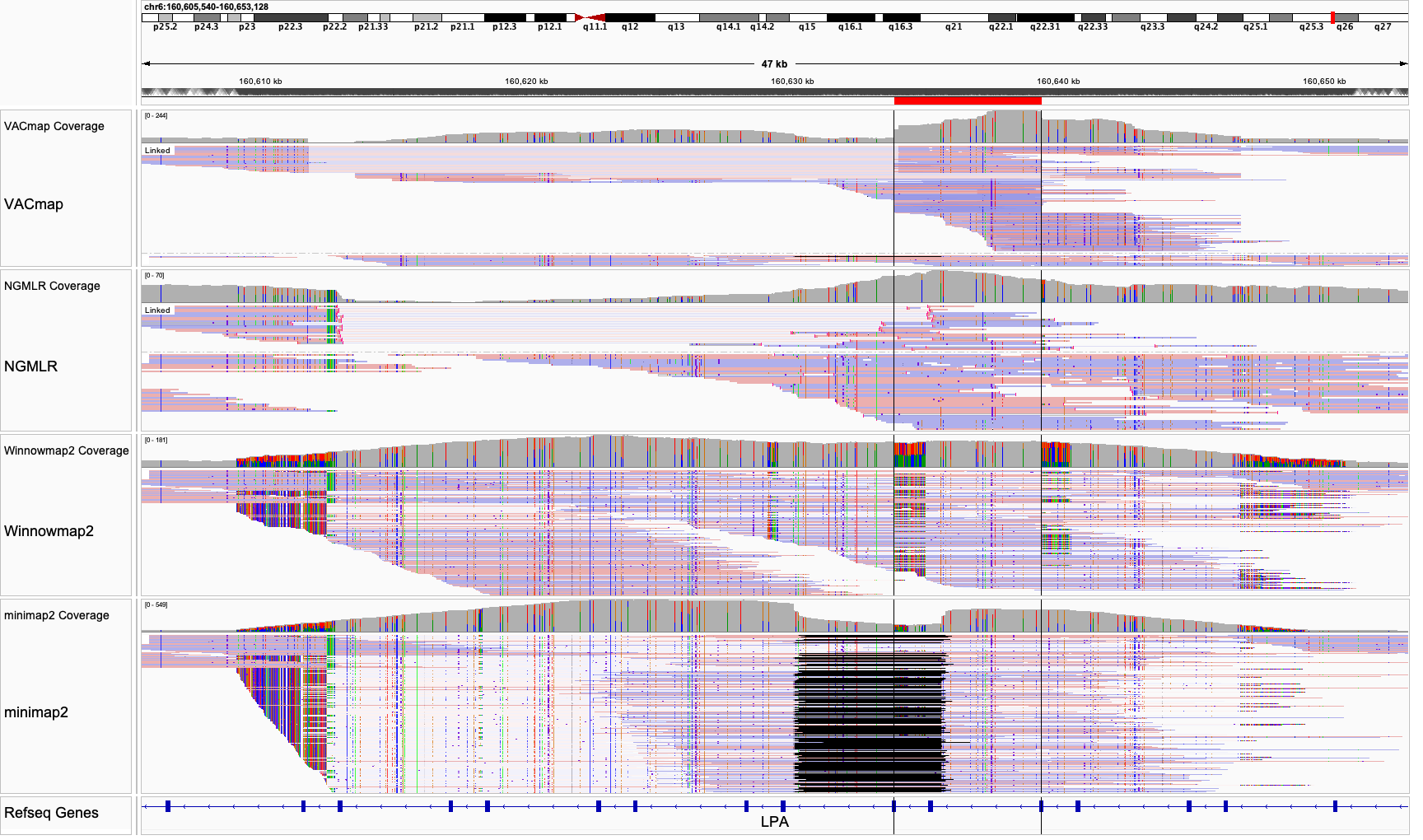

**Supplementary Figure 14 The IGV visualization of HG002 HiFi alignments produced by four aligners under GRCh38 reference in the KIV-2 region (GRCh38).**

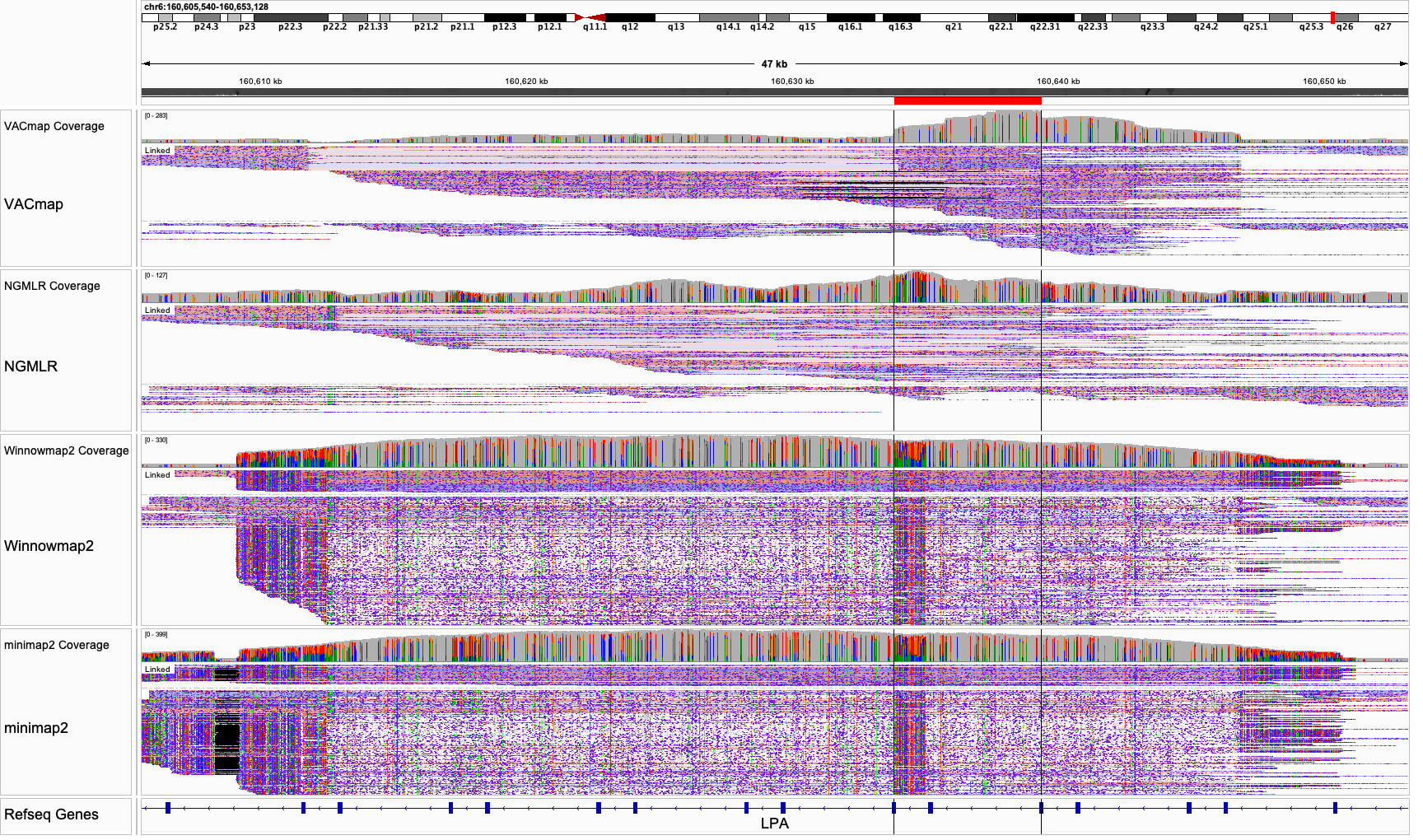

**Supplementary Figure 15 The IGV visualization of HG002 ONT alignments produced by four aligners under GRCh38 reference in the KIV-2 region (GRCh38).**

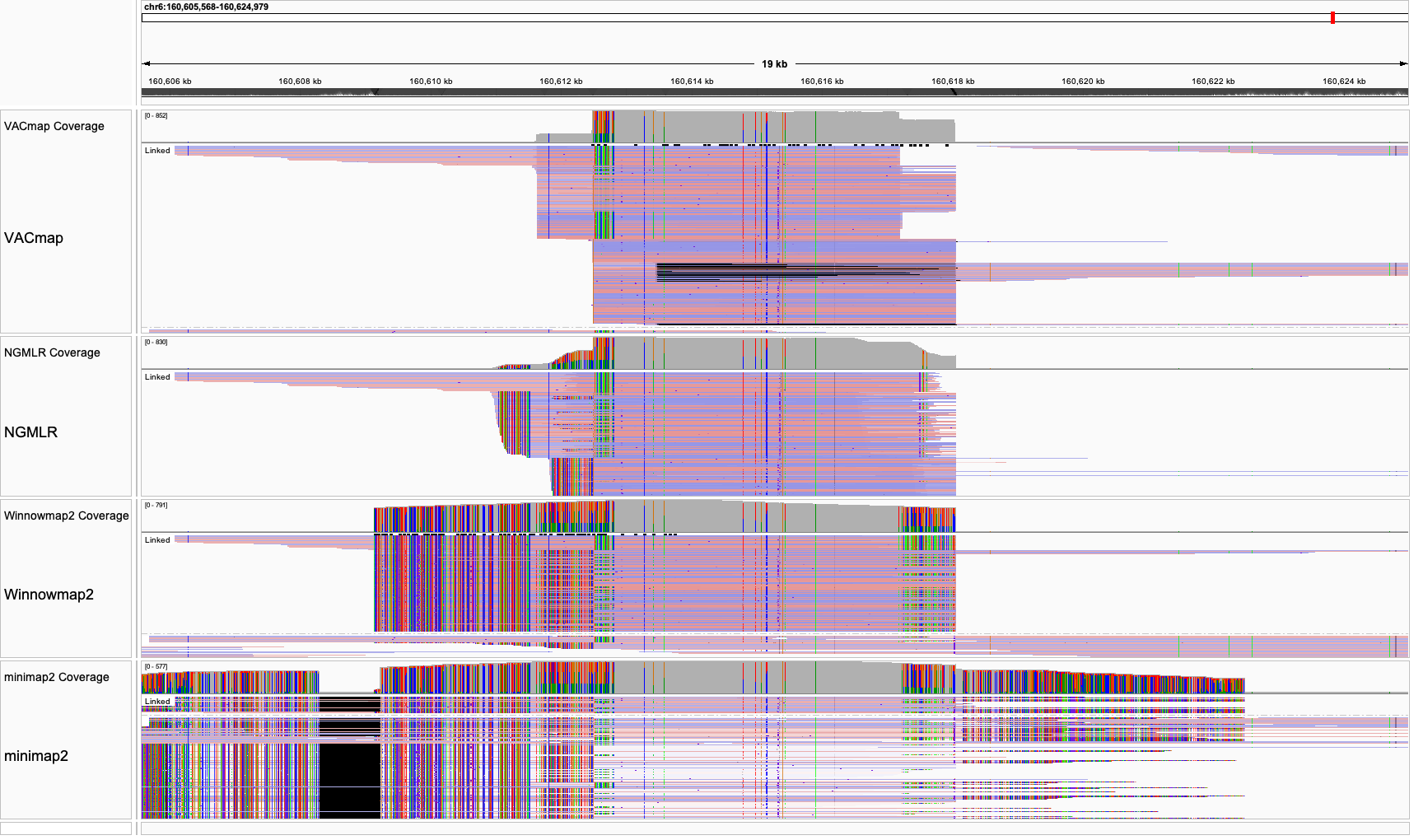

**Supplementary Figure 16 The IGV visualization of CHM13 HiFi alignments produced by four aligners under modified GRCh38 reference in the KIV-2 region.**

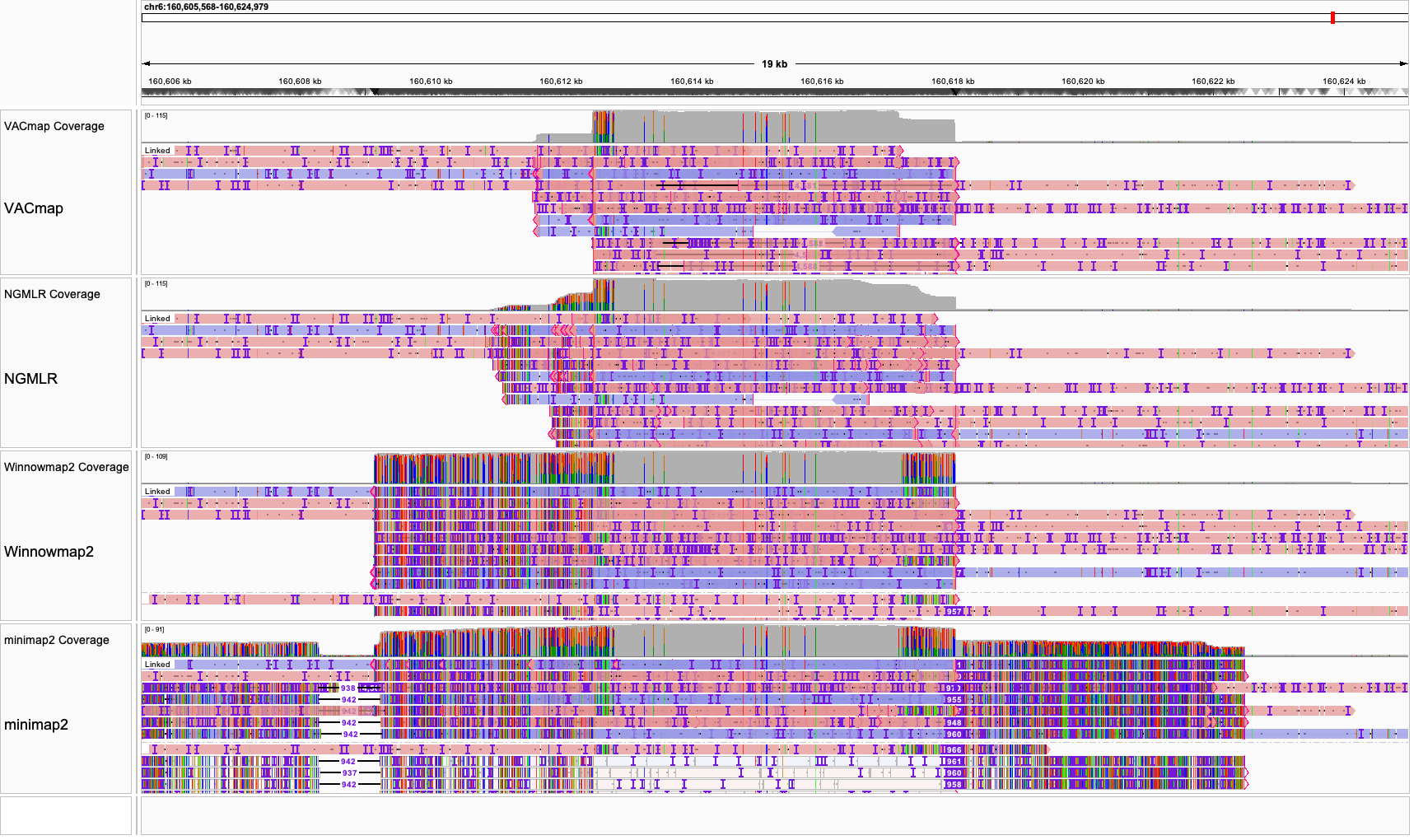

**Supplementary Figure 17** **The IGV visualization of CHM13 ONT alignments produced by four aligners under modified GRCh38 reference in the KIV-2 region.**

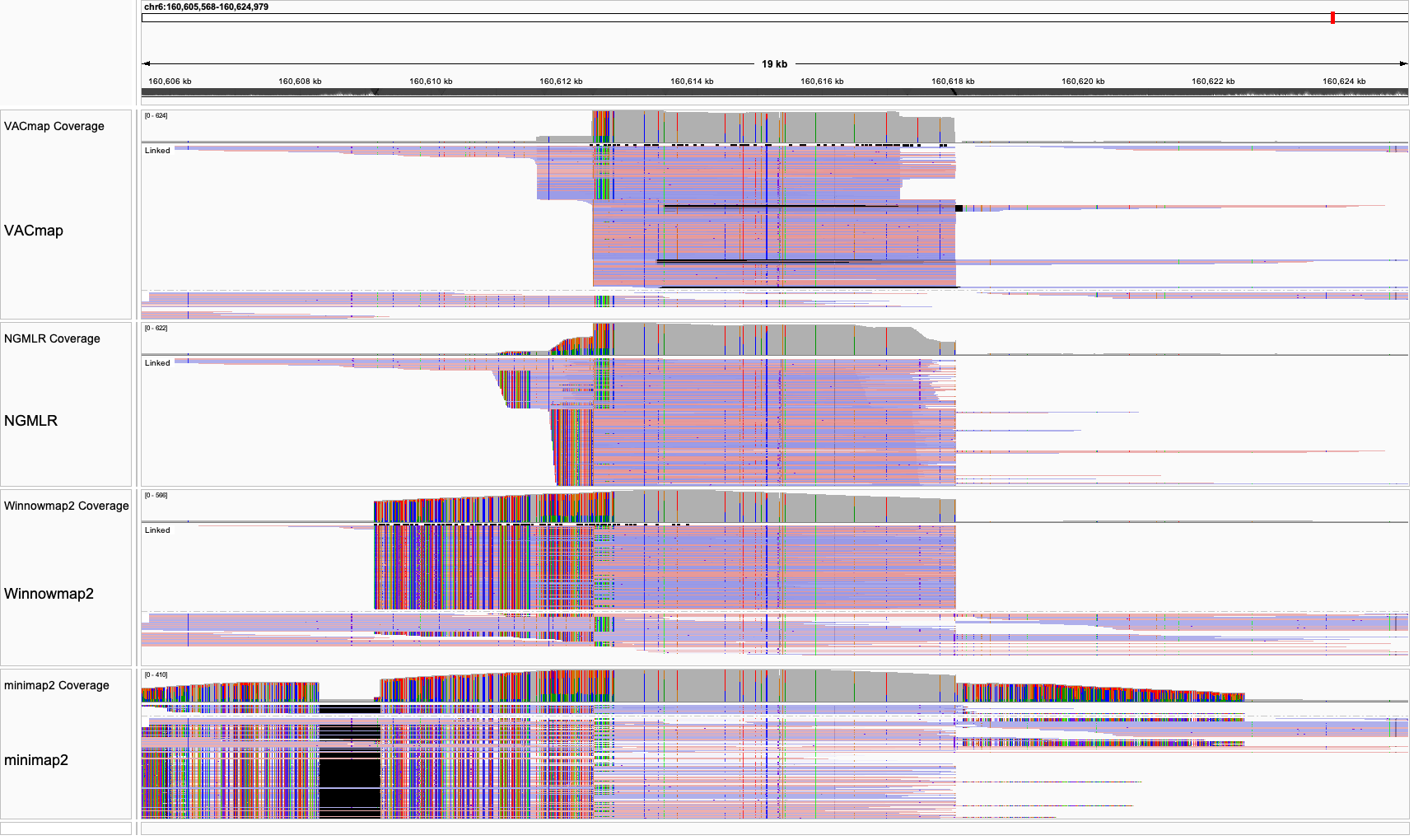

**Supplementary Figure 18** **The IGV visualization of HG002 HiFi alignments produced by four aligners under modified GRCh38 reference in the KIV-2 region.**

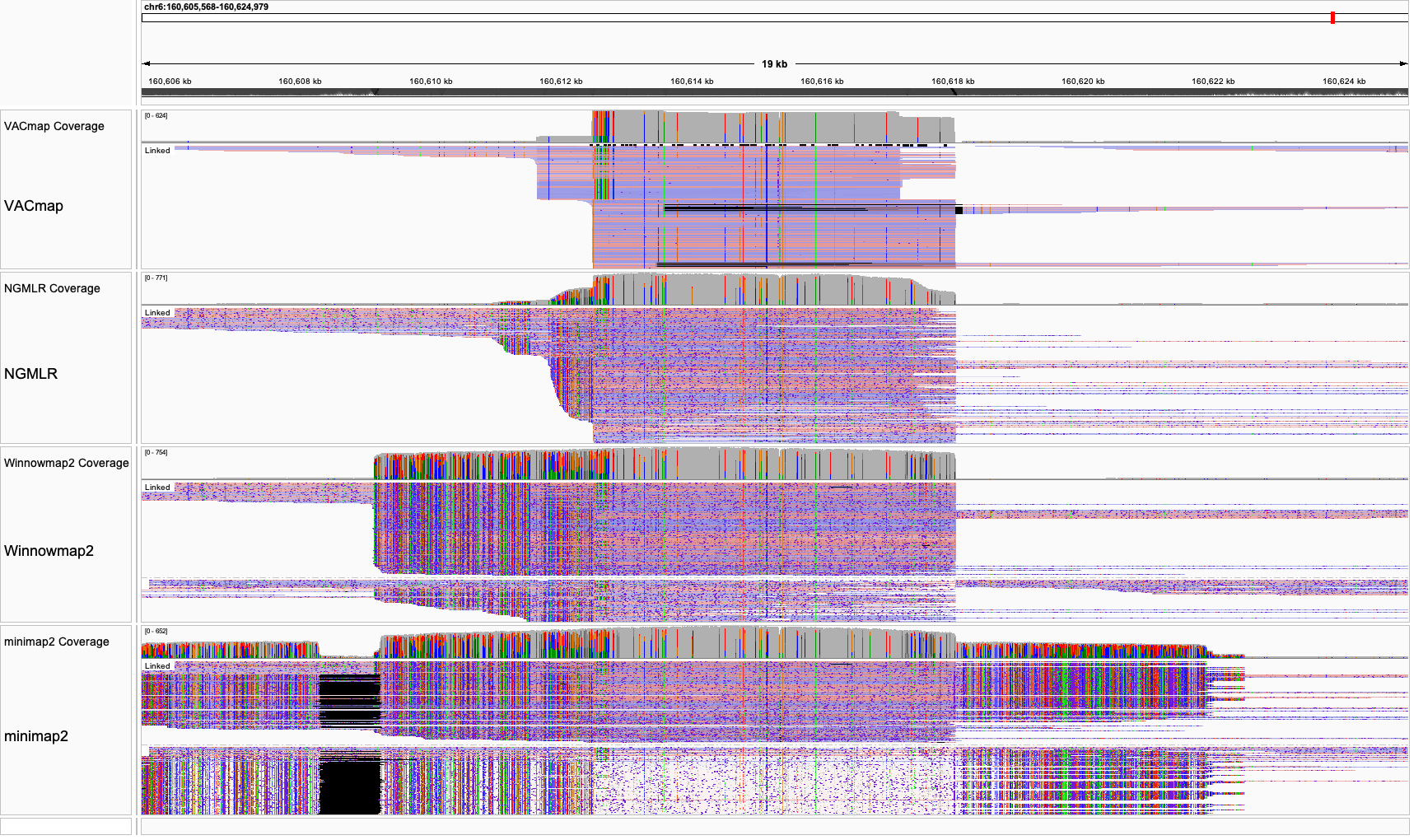

**Supplementary Figure 19 The IGV visualization of HG002 ONT alignments produced by four aligners under modified GRCh38 reference in the KIV-2 region.**

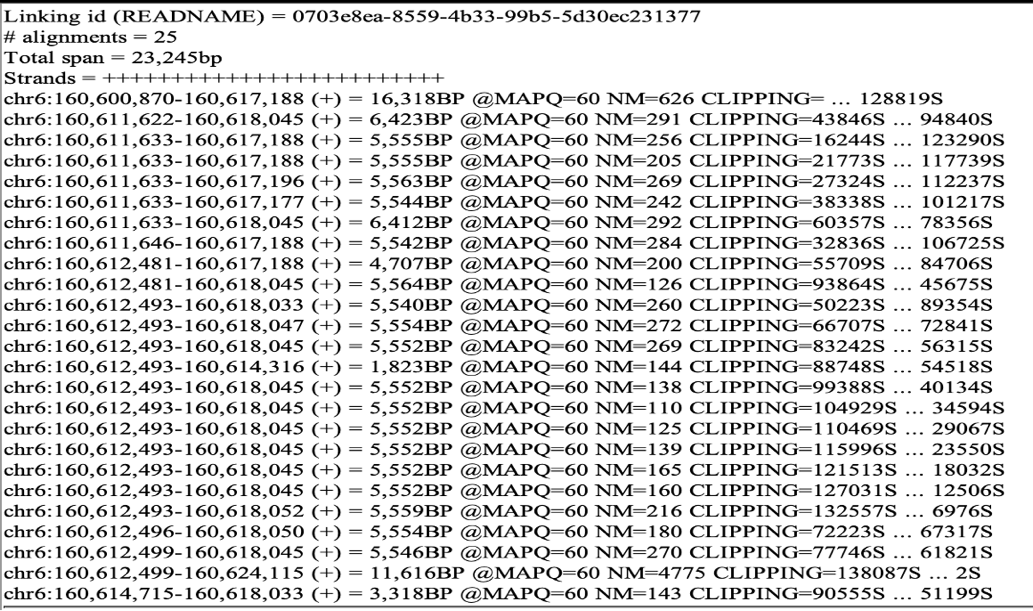

**Supplementary Figure 20 The alignment detail of an ONT long read from CHM13 human sample resolved all 23 KIV-2 repeat units.**

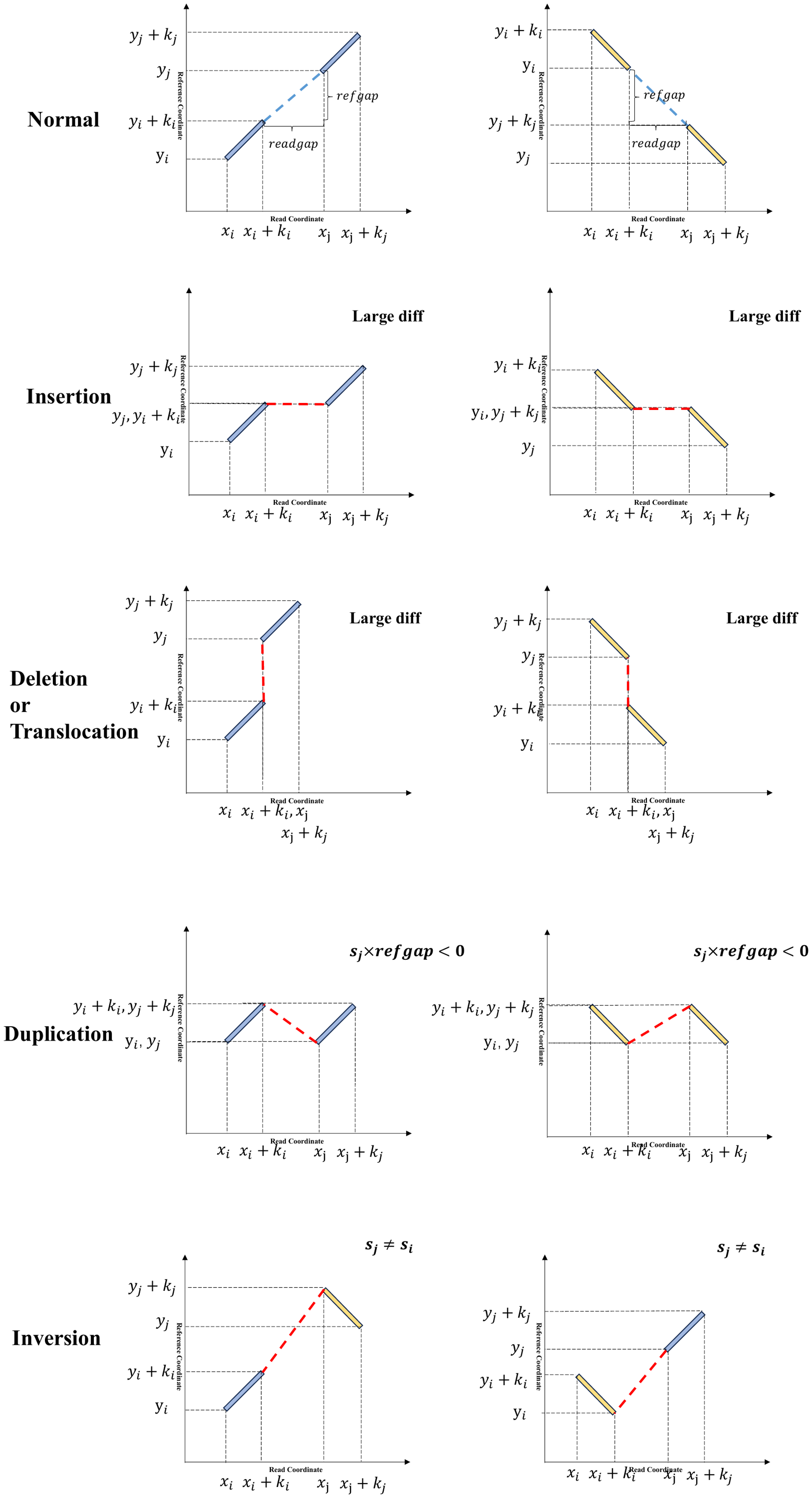

**Supplementary Figure 21 The Examples for assigning normal (blue) or variation (red) edges in different scenarios.**
